# Personality modulates brain responses to emotion in music: Comparing whole-brain and regions-of-variance approaches

**DOI:** 10.1101/651133

**Authors:** Kendra Oudyk, Iballa Burunat, Elvira Brattico, Petri Toiviainen

## Abstract

Whether and how personality traits explain the individual variance in neural responses to emotion in music remains unclear. The sparse studies on this topic report inconsistent findings. The present study extends previous work using regions of variance (ROVs) as regions of interest, compared with whole-brain analysis. Fifty-five subjects listened to happy, sad, and fearful music during functional Magnetic Resonance Imaging. Personality was measured with the Big Five Questionnaire. Results confirmed previous observations of Neuroticism being positively related to activation during sad music, in the left inferior parietal lobe. In an exploratory analysis, Openness was positively related to activation during Happy music in an extended cluster in auditory areas, primarily including portions of the left Heschl’s gyrus, superior and middle temporal gyri, supramarginal gyrus, and Rolandic operculum. In the whole-brain analysis, similar results were found for Neuroticism but not for Openness. In turn, we did not replicate previous findings of Extraversion associated to activity during happy music, nor Neuroticism during fearful music. These results support a trait-congruent link between personality and emotion-elicited brain activity, and further our understanding of the action-observation network during emotional music listening. This study also indicates the usefulness of the ROV method in individual-differences research.

## Introduction

In everyday life, people often describe each other in terms of characteristics or traits. In research, there is interest in whether biological differences underlie these patterns of behavior and cognition. Some individual differences, such as emotional tendencies, are relevant to both clinical and non-clinical contexts; healthy individuals vary in their emotional tendencies, but some tendencies may be related to mental health. In particular, the personality traits Extraversion and Neuroticism have been respectively related to behavioral tendencies to experience positive and negative emotions, both in laboratory settings (Larsen & Ketelaar, 1991) and in everyday life (Bolger & Schilling, 1991; David, Green, Martin, & Suls, 1997; Marco & Suls, 1993; Pavot, Diener, & Fujita, 1990). Further, these traits have been related to subjective well-being (Richard & Diener, 2009), happiness (Costa & McCrae, 1980), mood disorders (Jylhä & Isometsä, 2006), and emotional intelligence (Craig et al., 2009). Thus, research on the biological correlates of these traits may further our understanding of both normal variation between individuals as well as mental health outcomes.

Quantitative research on personality became more prominent in the later twentieth century, and with this research came theories about its biological basis, which often referred to the neural underpinnings of personality (e.g. Cloninger, 2000; Eysenck, 1967; Gray, 1984). In the past few decades, with advances in noninvasive brain imaging technology such as functional magnetic resonance imaging (fMRI), it became possible to investigate correlations between brain activity and individual differences like personality. A body of research has emerged investigating the relationship between different traits and neural activations to stimuli portraying different emotions. This research has shown that personality may be related to neural responses to emotional stimuli in a variety of limbic and multimodal regions of the brain (see the Supplementary Materials for details on this literature search, and see Table 1 for the findings of the studies found in this search).

**Table 1.**
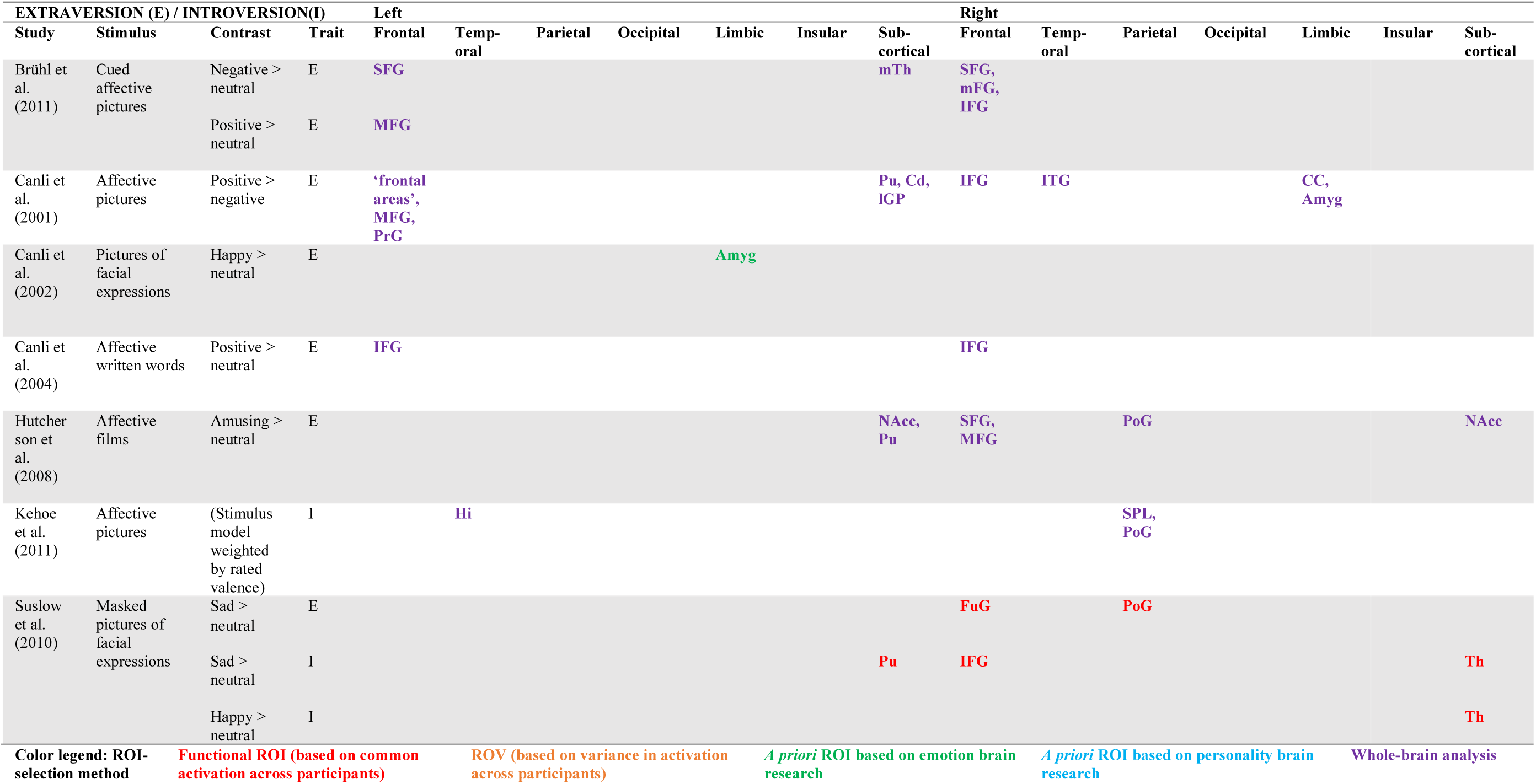
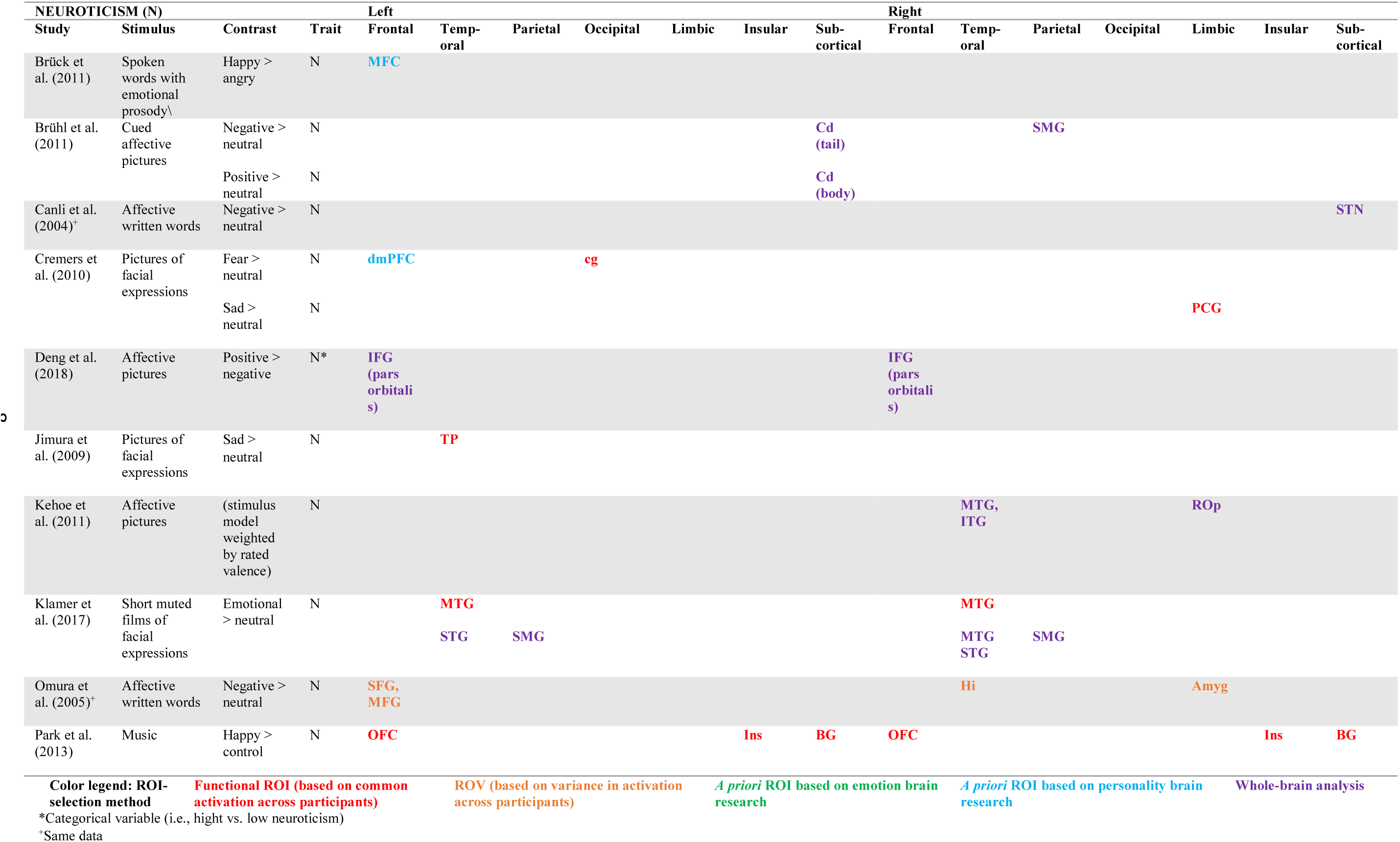
Significant results in studies on the role of Extraversion/Introversion and Neuroticism (next page) in neural activation to emotional stimuli. Amyg: amygdala; BG: basal ganglia; CC: cingulate cortex; Cd: caudate; Cd: caudate; cg: calcarine gyrus; dmPFC: dorsomedial prefrontal cortex; FuG: fusiform gyrus Hi: hippocampus; IFG: inferior frontal gyrus; IGP: inferior globus pallidus; Ins: insula; ITG: inferior temporal gyrus;ITG: inferior temporal gyrus; MFC: middle frontal cortex; mFG: medial frontal gyrus; MFG: middle frontal gyrus; MTG: middle temporal gyrus; MTG: middle temporal gyrus; mTh: medial thalamus; NAcc: nucleus accumbens; OFC: orbitofrontal cortex; pCG: posterior cingulate gyrus; PoG: postcentral gyrus; PrG: precentral gyrus; Pu: putamen; Rop: rolandic operculum; SFG: superior frontal gyrus; SMG: supramarginal gyrus; SPL: superior parietal lobe; STG: superior temporal gyrus; STN: subthalamic nucleus; Th: thalamus; TP: temporal pole.

Most of these studies used visual stimuli such as emotional facial expressions or words, with one study using sentences spoken with emotional prosody. However, the picture is less clear when it comes to music as an emotional stimulus. Based on the literature search performed for the present study (see the Supplementary Materials for literature search methods), there has only been one study on the role of personality in neural activations to emotion in music. This study, by Park et al. (2013), had some unexpected findings regarding trait-emotion pairings and a small sample size.

Outside the literature search, which was limited to studies reporting activations, we found one additional study investigating the role of personality in eigenvector centrality (a graph theoretical measure related to connectivity) during music listening (Koelsch, Skouras, & Jentschke, 2013); they found no significant results for Extraversion or for Neuroticism.

This lack of research on the role of personality in brain activity during perception of musical stimuli is noteworthy for several reasons. First, music is linked to the perception, recognition, and induction of emotion. In everyday contexts, music is used by listeners to experience and regulate their emotions (North, Hargreaves, & Hargreaves, 2004; Sloboda, O’Neill, & Ivaldi, 2001), and listeners report that music induces emotions more than half of the time that they are listening (Juslin & Laukka, 2004). There is evidence for cross-cultural consistency in the perception of emotion in music (e.g., Balkwill & Thompson, 1999; Balkwill, Thompson, & Matsunaga, 2004; Fritz et al., 2009, c.f., Jacoby & McDermott, 2017; McDermott, Schultz, Undurraga, & Godoy, 2016). Music can elicit core components of everyday emotions (Scherer, 2005), such as bodily symptoms and motor expressions (Lundqvist, Carlsson, Hilmersson, & Juslin, 2009), subjective feelings (Juslin & Laukka, 2004), and action tendencies (Burger, Saarikallio, Luck, Thompson, & Toiviainen, 2013). Further, music perception is related to neural activity in brain areas associated with processing emotion and reward (for a review, see Koelsch, 2014). Thus, music can be a useful tool for investigating the neural correlates of emotion (Koelsch, 2018).

Second, there is some behavioral evidence of a link between personality and emotion in music. Extraversion, a trait characterized by positive-emotion tendencies (Costa Jr & McCrae, 1992), has been related to preference for musical genres that “emphasize positive emotions and are structurally simple” (p. 1241; Rentfrow & Gosling, 2003) and those that “are lively and often emphasize rhythm” (p. 1242). Further, Extraversion has been related to liking happy-sounding music (Vuoskoski & Eerola, 2011b), and higher felt emotions during music listening (Vuoskoski & Eerola, 2011a). On the other hand, Neuroticism, characterized by negative-emotion tendencies (Costa Jr & McCrae, 1992), has been related to higher felt sadness while listening to music (Ladinig & Schellenberg, 2012, c.f. Vuoskoski & Eerola, 2011a, 2011b). When music is being used as an emotional stimulus, the trait Openness to Experience might modulate the emotional responses, since music is an art form in addition to being emotional, and Openness is related to aesthetic sensitivity (Costa Jr & McCrae, 1992). Indeed, it has been related to to experiences of awe (Silvia, Fayn, Nusbaum, & Beaty, 2015) and chills (Colver & El-Alayli, 2016) while listening to music. Regarding emotion, Openness has been associated with greater liking for sad music (Vuoskoski & Eerola, 2011b; Vuoskoski, Thompson, McIlwain, & Eerola, 2012), fearful music (Vuoskoski & Eerola, 2011b), and sadness-inducing music (Ladinig & Schellenberg, 2012), and to the intensity of emotions induced by sad and tender music (Vuoskoski & Eerola, 2011a).

This behavioral research indicates that music is an emotional stimulus and that personality may be related to experiences of emotion in music, warranting an investigation of potential biological underpinnings. But while the universal neural correlates (common to the general population) of music-evoked emotions has been well characterized (for a review, see Koelsch, 2014), it remains unclear whether emotional tendencies modulate the individual variations in emotional responses to music observed in a variety of behavioral studies and in their underlying neural correlates. Given the lower effect sizes in individual differences research (Gignac & Szodorai, 2016), greater statistical power may be needed to answer these questions. To increase the power, the methods of the previous study on the role of personality in neural activations to emotion in music (Park et al., 2013) may be extended by using a larger sample size (from 12 to 55) and a different method for selecting regions of interest (ROIs).

Park et al. (2013) selected ROIs based an initial analysis of regions consistently activated across participants. However, regions with consistent activation across participants may not be ideal candidate regions for investigating individual differences between participants (Omura, Aron, & Canli, 2005). Arguably the least-biased approach regarding brain areas would be to perform a whole-brain analysis. However, this analysis has a higher risk of Type II errors; correcting for multiple comparisons over all voxels in the brain may make the analysis too stringent to find true results, especially in this case where the effect size is likely small (Gignac & Szodorai, 2016). In order to have better statistical power than could be had in a whole-brain analysis, and at the same time avoid limiting functional ROIs to regions of consistent activation across participants, Omura et al. (2005) proposed using regions of higher between-subjects variance (relative to within-subjects variance) as ROIs for investigating individual differences (Omura et al., 2005). The present study further builds on the previous study by Park et al. (2013) by using this alternative, and perhaps more effective, method for selecting ROIs as regions of variance (ROVs). A whole-brain analysis is additionally included as a benchmark against which the results of ROV analysis can be compared.

Thus, the goal of the present study is to further investigate the role of personality in brain activations to emotion in music, using the regions-of-variance method in comparison to whole-brain analysis. We hypothesized that Extraversion would be related to activations to happy music and Neuroticism to sad and/or fearful music, given their characteristic emotional tendencies. The investigation of Openness to Experience was exploratory, since it is not characterized by emotional tendencies. Regarding brain areas, it was expected that these personality variables would be related to emotion-related activation in limbic and multimodal association areas (note that these expectations were not explicitly tested as hypotheses, since a data-driven method was used for selecting ROIs). Further, we hypothesized that the ROV analysis would be more sensitive to extract individual differences in personality associated with affective music listening, compared to the whole-brain analysis. Hence, we expect to obtain significant findings mainly with ROV and less with whole-brain analysis.

## Method

The paradigm used in this study was part of a larger experimental session (Bogert et al., 2016; Carlson et al., 2015), which belonged to the Tunteet protocol. It received ethics approval from the Koordinoiva ethical committee of Helsinki and Uusimaa Hospital District, and it was run according to the ethical guidelines of the Declaration of Helsinki.

### Participants

Sixty-three subjects were recruited via email lists from the University of Helsinki (Helsinki, Finland) and Aalto University (Espoo, Finland). Potential participants were excluded if they had hearing or neurological problems, or if they were taking psychopharmacological medicine. Prior to scanning, participants were informed about the study, consented to participate, and completed an fMRI safety questionnaire. They were compensated for every half-hour of participation with a voucher for cultural and exercise activities.

Following data collection, eight subjects were excluded from further analyses for the following reasons: too much movement (3), neuroradiological abnormalities diagnosed by a radiologist (3), technical issues (1), and incomplete personality data (1).

Thus, 55 participants (20-53 years, *M* = 28.6, *SD* = 7.98; 23 males; 3 left-handed) were included the final sample.

### Stimuli

The stimuli consisted of 30, 4-second instrumental music excerpts representing happiness, sadness, and fear (10 clips per emotion). To select these stimuli, 81 excerpts were taken from a film-music database with validated emotional labels (Eerola & Vuoskoski, 2011). Each excerpt was rated on nine emotions (happiness, sadness, fear, anger, tenderness, pleasure, disgust, surprise, unclear, or other emotion) by ten subjects who did not participate in the brain imaging study. The ten most representative excerpts for each Happy, Sad, and Fearful were selected as stimuli (30 total).

Adobe Audition was used to edit the stimuli so that they had equal loudness (average root mean square) and so that they lasted four seconds with 500 ms of fade-in and a fade-out. For more details on the stimuli, see Bogert et al. (2016).

### Procedure

Participants were presented with the question “How many instruments do you hear?” for 10 seconds, followed by a 4-second music clip and then a 5-second answer period in which they selected “one”, “two”, or “many” (see Figure 1). There was also an explicit emotion-labeling block during the same scanning session; only data from the implicit block was analyzed in the present study since it was more powerful to activate emotion-related limbic regions as compared to the other block, for the sake of maximizing statistical power toward the study of individual differences in emotional brain responses. In order to familiarize the participants with the tasks, there was a short training session before the study stimuli were presented.

**Figure 1:**
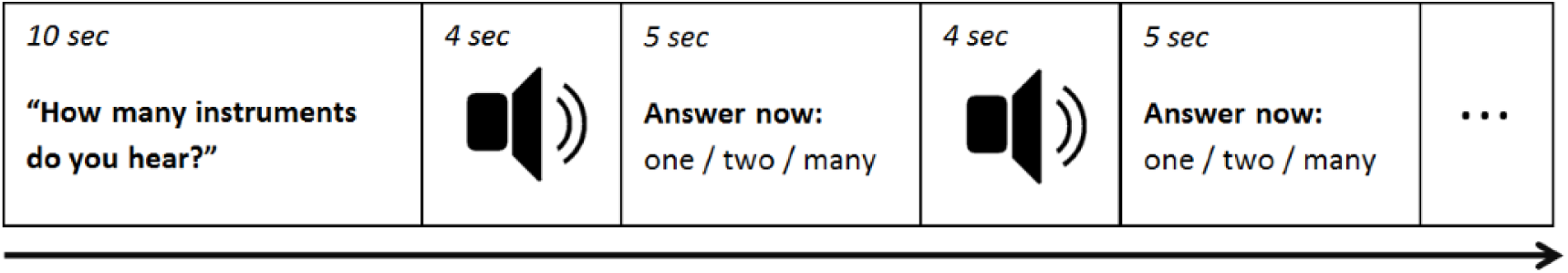
Stimulus presentation paradigm in the implicit emotion-processing block.

Personality was measured using the Big Five Questionnaire (BFQ; John & Srivastava, 1999) a separate day in the Biomag laboratory of the Helsinki University Central Hospital, along with other questionnaires and physiological measures. Among these additional questionnaires was the NEO-FFI (Costa & McCrea, 1992), another inventory of the Five-Factor Model of personality. However, the BFQ was used in further analysis because it had higher inter-subject reliability.

### fMRI acquisition

The apparatus used was a 3-T MAGNETOM Skyra whole-body scanner and a stan dard 20-channel head-neck coil (Siemens Healthcare, Erlangen, Germany), located in the Advanced Magnetic Imaging (AMI) Center at Aalto University, Espoo, Finland.

The functional images were obtained with interleaved gradient echoplanar imaging (EPI) with a 2-second repetition time (TR) and a 2-millisecond echo time. Thirty-three 4-mm slices were taken at a flip angle of 75°, resulting in 3 x 3 x 4 mm voxels with 0 mm spacing. The field of view was 192 x 192 mm, and the matrix 64 x 64 mm.

The structural images were taken following the fMRI task; they consisted of high-resolution anatomical T1-weighted MR images, with 176 slices, a 256 mm field of view, a 256 x 256 matrix, and 1 mm^3^ voxels with 0 mm spacing.

### fMRI preprocessing

Functional images were corrected for between-scan movement and slice timing, resampled to a 2-mm isotropic voxel size, spatially normalized to the MNI template using a 12-parameter affine transformation, and smoothed with a 6-mm FWHM Gaussian filter. Images were then examined on criteria for high quality and scan stability (*<*2 mm translation and *<*2*^o^* rotation during small-motion correction). These preprocessing steps were carried out in the SPM8 toolbox in MATLAB. The six parameters of head movement were entered as predictors of the BOLD signal in a multiple linear regression, and the residual was retained for further analysis. Long-term trends were removed using spline interpolation with 5 anchor points. The data were mean-centered within voxels. These temporal preprocessing steps were done in MATLAB using custom code.

### fMRI analysis

Data were analyzed using a voxelwise 2-level summary statistics approach (Holmes & Friston, 1998). In the first level, the BOLD signal for each participant was related to the condition regressors in a fixed-effects model using multiple linear regression. The estimated parameter weights for each emotional condition were contrasted against the other two emotion conditions, and then related to the personality data using a random-effects model (Holmes & Friston, 1998) using partial correlation. The sampling distribution of the resulting correlation coefficients was normalized with Fisher’s transformation (Fisher, 1915). For each emotion condition, a ROV map of F-statistics was calculated (Omura et al., 2005) and thresholded using the change point in the sorted list of F-values. This gave binary ROV masks that were then applied to the level-2 Z-maps. The ROVs for each emotional condition can be seen in Figure S3. A whole-brain analysis was also carried out for comparison with the ROV results. For both the ROV and whole-brain analyses, correction for multiple comparisons was done using randomization tests (Poldrack, 2007). In order to find cluster-size and -mass thresholds that corrected for multiple comparisons at a family-wise error rate (FWER) of 5%, Monte Carlo simulations with 10000 permutations of participant labels were carried out on the Z-maps (Nichols & Holmes, 2002). See the Supplementary Materials for a detailed description of these analysis steps and a note on avoiding circular analysis in functional ROI selection. These analyses were carried out in MATLAB using custom codes.

Data were analyzed using a voxelwise 2-level summary statistics approach (Holmes & Friston, 1998). In the first level, the BOLD signal for each participant was related to the condition regressors in a fixed-effects model using multiple linear regression. The estimated parameter weights for each emotional condition were contrasted against the other two emotion conditions (e.g., [2*happy] ¿ [sad + angry]), and then related to the personality data using a random-effects model (Holmes & Friston, 1998) using partial correlations to investigate each of the personality traits while controlling for the other two. The sampling distribution of the resulting correlation coefficients was normalized with Fishers transformation (Fisher, 1915). For the ROV analysis, a map of F-statistics was calculated for each emotional contrast (Omura et al., 2005) and thresholded using the change point in the sorted list of F-statistics; this gave binary ROV masks that were then applied to the level-2 Z-maps. Statistical inference and correction for multiple comparisons was performed using separate randomization tests for the ROV and whole-brain analyses (Poldrack, 2007). In order to find cluster-size and -mass thresholds that corrected for multiple comparisons at a family-wise error rate (FWER) of 5%, Monte Carlo simulations with 10000 permutations of participant labels were carried out on the Z-maps (Nichols & Holmes, 2002). See the Supplementary Materials for a detailed description of these analysis steps, for images of the ROV maps, and for a note on avoiding circular analysis in functional ROI selection. These analyses were carried out in MATLAB using custom code.

## Results

Neuroticism was positively related to activation during Sad music in the left inferior parietal lobe, including the supramarginal gyrus (BA 40) and angular gyrus (BA 39; cluster-defining threshold, CDT, of *p* = 0.001, corrected at 5% FWER). The whole-brain analysis revealed a larger cluster in a similar location, additionally with voxels in the postcentral gyrus (BA 2). All voxels in the cluster found with the ROV analysis were included in the cluster found with the whole-brain analysis. See Table 2 for details and Figure 2 for orthographic views of the statistical maps of these results.

**Figure 2.**
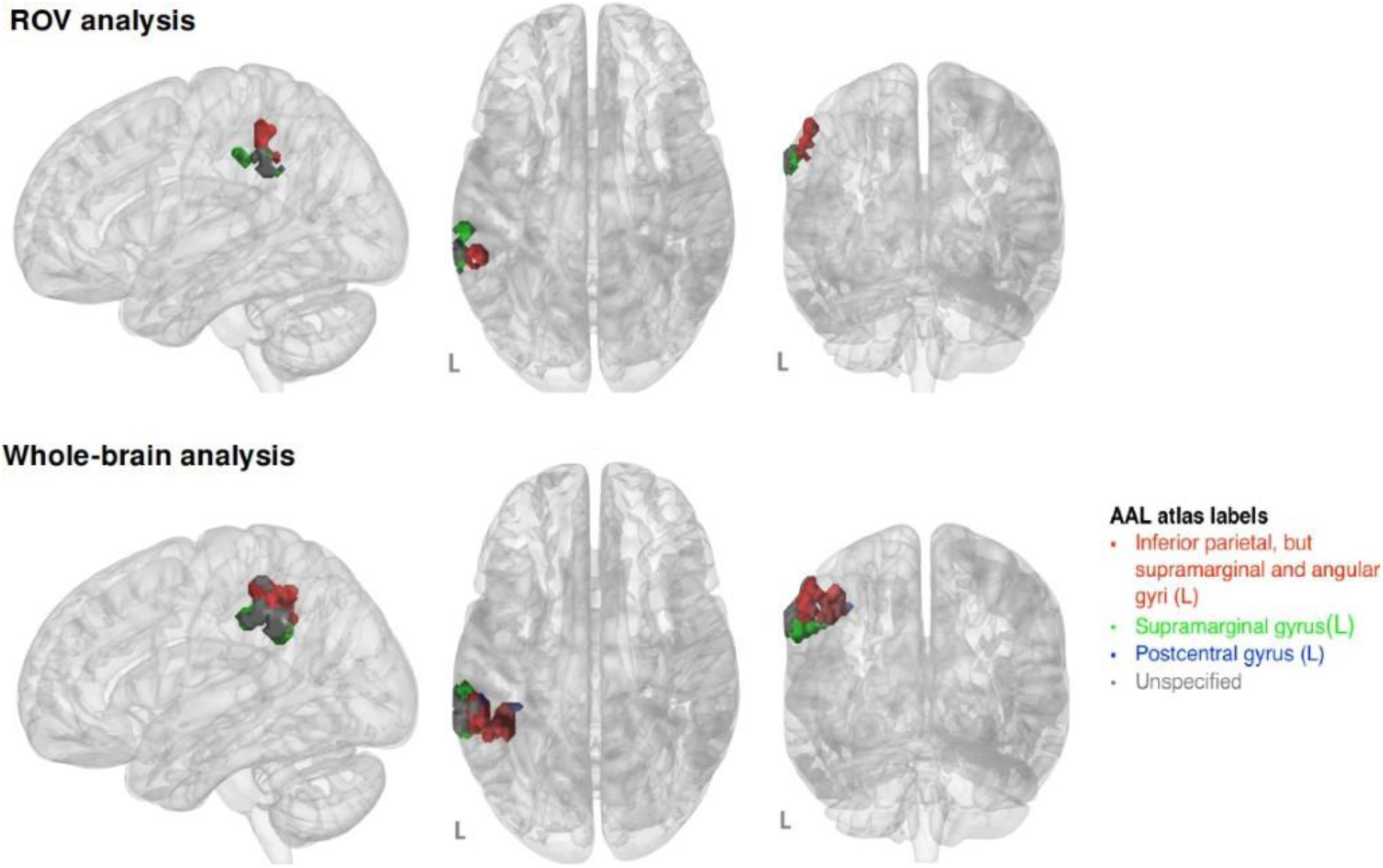
Significant cluster for Neuroticism & Sad music. The ROV result was found with a CDT of *p* = 0.001, corrected at 5% FWER using non-parametric cluster-size and -mass thresholds, which were *k* = 68 voxels and Z = 248.49, respectively. The whole-brain result was found with a CDT of *p* = 0.001, corrected at 5% FWER using non-parametric cluster-size and -mass thresholds, which were *k* = 207 voxels and Z = 760.68. All voxels in the ROV results were included in the whole-brain results.

**Table 2:**
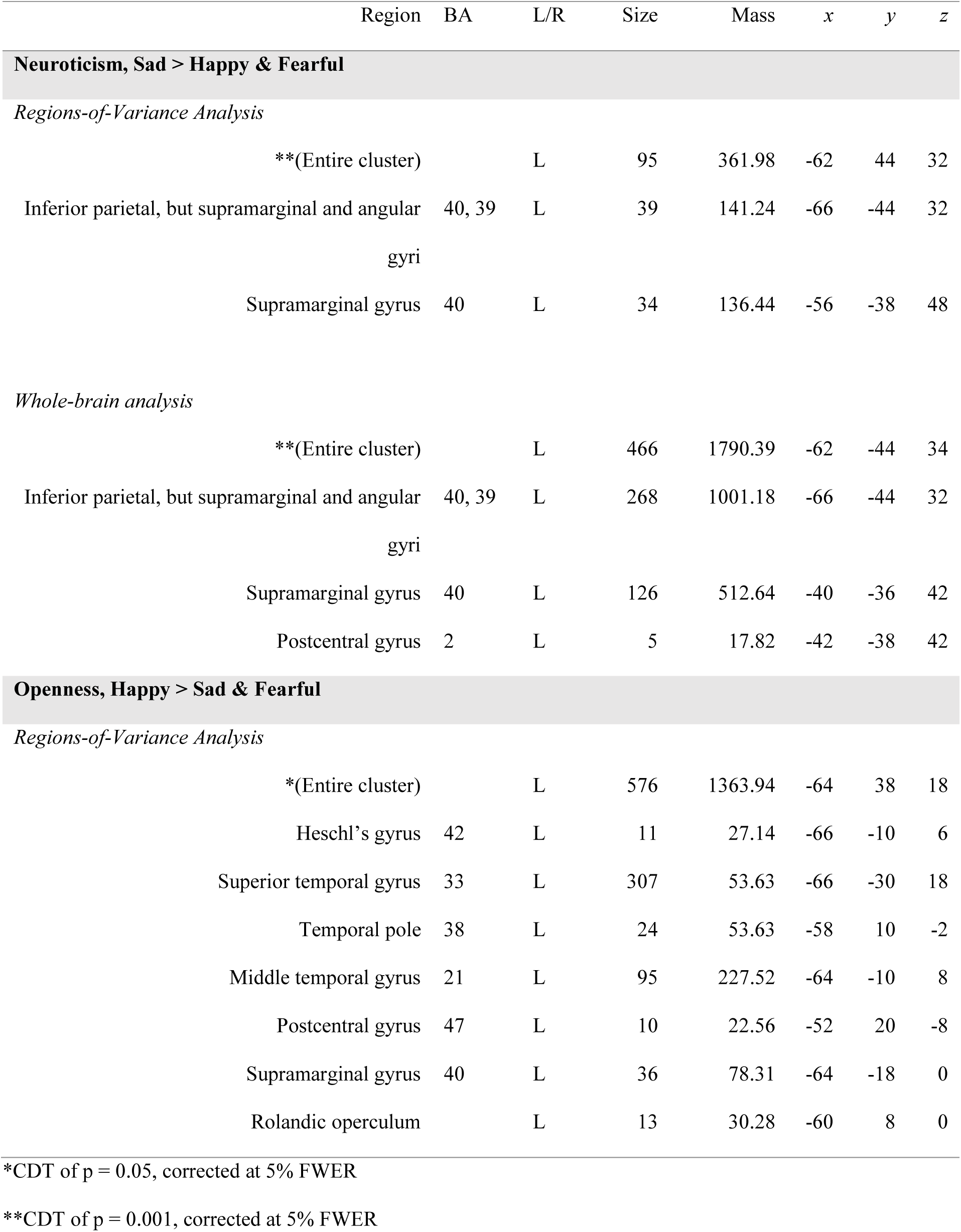
Significant results. All results passed correction for multiple comparisons at a 5% FWER, using cluster-size and cluster-mass thresholds determined in a nonparametric permutation test for each combination of trait and emotion contrast. Thus, entire clusters were accepted/rejected in an omnibus test, and it should be noted that while the component regions and their peak-value locations are listed in this table for the purpose of illustrating cluster extent, corrected significance was not estimated for individual voxels. Region = region label, from the Automatic Anatomic Labeling (AAL) atlas. BA: Brodmann area corresponding to the region. L/R: left/right hemisphere. Size: number of voxels in the cluster, each being 8 mm^3^ isotropic. Mass: mass of the cluster, which is the sum of the Z-values. *x, y, z*: MNI coordinates of the peak voxel in the given region.

Openness was significantly and positively related to activation during Happy music in an extended cluster in the region of the left superior temporal lobe, and primarily included voxels in Heschl’s gyrus (BA 42) and the superior temporal gyrus (BA 22), extending into the temporopolar area (BA 38), middle temporal gyrus (BA 21), postcentral gyrus (BA 2), supramarginal gyrus (BA 40), and the Rolandic operculum. This cluster was significant with a CDT of *p* = 0.05, corrected at 5% FWER. This cluster was not significant in the whole-brain analysis, with the more stringent FWE-corrected cluster-size threshold. See Table 2 for details and Figure 3 for orthographic views of the statistical map of this result.

**Figure 3.**
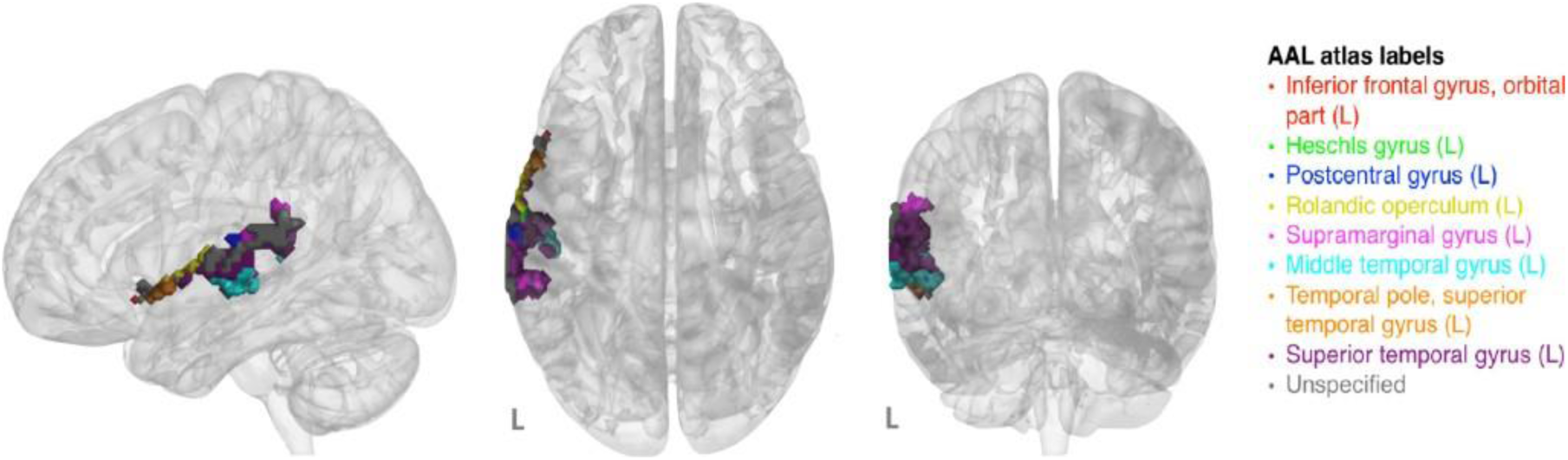
Significant cluster for Openness & Happy music (ROV analysis). This result was found with a CDT of *p* = 0.05, corrected at 5% FWER using non-parametric cluster-size and -mass thresholds, which were *k* = 507 voxels and Z = 1295.28, respectively.

The results were the same after cluster-size and -mass thresholding, suggesting that for this dataset, cluster size and mass are monotonically related at the high end of their distributions. There were no results that passed correction for multiple comparisons for any other combination of trait and emotional condition.

## Discussion

This study investigated the role of personality in the neural correlates of perceiving emotions in music. Specifically, it looked at activations associated with Extraversion, Neuroticism, and Openness to Experience during implicit perception of Happy, Sad, and Fearful emotions portrayed by instrumental music.

Extraversion and Neuroticism are characterized by tendencies to experience positive and negative emotional states (e.g., Larsen & Ketelaar, 1991, and have been related to brain function during perception of positive and negative emotional stimuli, respectively (e.g., Canli, Sivers, Whitfield, Gotlib, & Gabrieli, 2002; Canli et al., 2001; Haas, Constable, & Canli, 2008). Thus, it was expected that Extraversion would be related to the hemodynamic activity during perception of Happy music, and Neuroticism during Sad or Fearful music. There were no significant clusters that passed the correction for multiple comparisons for Extraversion & Happy or for Neuroticism & Fearful, but there was one robustly significant cluster in the left inferior parietal lobe for Neuroticism & Sad.

The inclusion of Openness to Experience as a personality trait of interest was exploratory, since Openness is not normally characterized by emotional tendencies. Openness was positively related to activation during Happy music, in a cluster including auditory association cortex, temporopolar area, medial temporal lobe, and some extension into the middle temporal gyrus, and postcentral gyrus.

### Neuroticism & sad music

Neuroticism was positively related to activation in a cluster in the left inferior parietal lobe (IPL). This was a robust finding; a similar cluster was found in both the ROV analysis and the whole-brain analysis, in both cases at the most conservative CDT.

There has been some research relating Neuroticism (or related traits) to function of the IPL. Neuroticism has been associated with connectivity between the retrosplenial cingulate cortex and the right IPL during reading of worrisome versus neutral sentences (Servaas, Riese, Ormel, & Aleman, 2014). Sander et al. (2005) found that Behavioral Inhibition System scores (BIS, measuring harm-avoidance motivation) were positively correlated with activity in the right IPL during perception of angrily-spoken nonsense words, and that Behavioral Activating System scores (BAS, measuring reward-approach motivation) scores were positively correlated with activity in the left IPL. This is relevant because BIS scores are positively related to Neuroticism (Jorm et al., 1998). It is notable that the present findings were in the *left* IPL, however, there has been mixed evidence concerning lateralization in research on BIS-BAS scores and brain function (Coan & Allen, 2003; Harmon-Jones & Allen, 1997).

The IPL has many functions, but notable for this context is its association with cognitive emotional processes. The temporoparietal junction (TPJ) has been related to cognitive empathy (the understanding of others’ emotional states; Schnell, Bluschke, Konradt, & Walter, 2011) and reasoning about others’ minds and beliefs (Samson, Apperly, Chiavarino, & Humphreys, 2004; Saxe & Kanwisher, 2003). The left IPL has been associated with suppression as a method of negative-emotion regulation (Goldin, McRae, Ramel, & Gross, 2008). This may be relevant here because sadness is a negatively-valenced emotion, but some research has suggested that Neuroticism is not related to suppression as an emotion-regulation strategy, but rather to reappraisal (Gross & John, 2003; Wang, Shi, & Li, 2009).

The IPL has also been associated with higher-level auditory perceptions, consistent with its activation here to music. The left IPL has been associated with phonological processing (Vallar & Papagno, 1995; Zatorre, Evans, Meyer, & Gjedde, 1992), processing of musical timbre (Alluri et al., 2012), musical performance and imagery (Meister et al., 2004), and perception of melodies versus random tones (BA 40) and of harmonized versus unharmonized melodies in musicians (BA 39; Schmithorst & Holland, 2003). In a study that used data from the same larger dataset as the present study, Bogert et al. (2016) looked at common activation across participants and found the bilateral IPL to be active during sad music.

Further, the left IPL is part of the action-observation network and putative mirror neuron system. The peak value for the action-observation network, as found in the meta-analysis by Caspers, Zilles, Laird, and Eickhoff (2010), was located near the cluster for Neuroticism and sad music found here (see Figure S7). Since this meta-analysis only reported the coordinates of the peak, and not the full extent of the left-IPL cluster, we also looked at the intersection between our results and the “action observation” term-based meta-analytic association test map from NeuroSynth, and found that the clusters did indeed overlap (see Figure S8). This is relevant because it has been proposed that emotional contagion (Juslin & Västfjäll, 2008) and empathy (Molnar-Szakacs & Overy, 2006) are central neural mechanisms behind emotional responses to music.

Thus, although the IPL is not part of networks traditionally associated with the experience of emotion, it has been related to emotional cognitive processes, processing of pitch and speech sounds associated with emotion, processing musical emotion, and it is part of the action-observation network.

### Openness & happy music

Openness was positively related to activation in an extended cluster in the area of the left superior temporal lobe, primarily including voxels in Heschl’s gyrus, the superior temporal gyrus, and the medial temporal lobe, with some extension into the ventral postcentral gyrus, posterior superior temporopolar area, middle temporal gyrus, and the Rolandic operculum. Although there appears to be little functional brain research on Openness with emotional stimuli, this finding is interesting in light of previous research on music and emotion.

Regarding music and emotion, the left superior temporal gyrus has been related to activation during happy music (Brattico et al., 2011, 2016; Concina, Renna, Grosso, & Sacchetti, 2019; Mitterschiffthaler, Fu, Dalton, Andrew, & Williams, 2007), and to disliked music (Brattico et al., 2016). Koelsch, Fritz, v. Cramon, Müller, and Friederici (2006) reported that perception of pleasant music is related to activity in the left Rolandic operculum as well as the *right* temporal pole, but activity in the left temporal pole did not reach significance. In a recent study, Koelsch, Skouras, and Lohmann (2018) found greater activation in the bilateral auditory cortex during joyful music, compared to fearful music, and they found functional connectivity between the auditory cortex and limbic, paralimbic, and neocortical areas. We might speculate that the auditory cortices include neural representations of auditory emotions, and that personality traits can implicitly modulate these auditory-cortex responses.

Emotion processing in other domains has been associated with function of regions in this cluster. The left temporal pole has been linked to complex perceptions with emotional responses (Olson, Plotzker, & Ezzyat, 2007) and to judgments of affective semantics (Ethofer et al., 2006), and the left Rolandic operculum has been related to perceptions of emotion in speech (Kotz et al., 2003).

Thus, the regions related to Openness & Happy in the present study have previously been associated with emotional processing, both in music and other modalities. In order to elucidate the brain correlates of role of Openness in perceiving music, future research may benefit by using longer musical stimuli to allow for higher engagement with the music and aesthetic responses, as well as physiological and/or verbal measures of engagement and emotional experience.

It should be noted that this finding was less robust, being found only in the ROV analysis and not in the whole-brain analysis, and only at the least-conservative CDT (*p* = 0.05). Eklund, Nichols, and Knutsson (2016) raised concern over the use of low CDTs, as they resulted in inflated empirical family-wise error rates when parametric methods for multiple-comparison methods were used. But in the case of the non-parametric method, which was used here, the FWERs were nominal even at a lower CDT (*p* = 0.01). That being said, they did not examine the empirical FWERs for a CDT of *p* = 0.05, so these results should still be interpreted cautiously.

### Null results

Contrary to the hypotheses, Extraversion was not related to activations during Happy music, nor was Neuroticism related to activations during Fearful music. These hypotheses were based on behavioral associations between these traits and positive and negative emotionality, respectively, and on trait-congruent associations between neural activity and personality during perception of emotional stimuli.

The null results in the present study may be due to a failure to detect an effect or to the absence of an effect. Associations between personality traits and brain activity during perception of emotional stimuli may have smaller effect sizes than mean activations (across participants) to emotional stimuli (Gignac & Szodorai, 2016), and thus be difficult to detect. Although there have been past associations between Extraversion and activation during perception of positive stimuli (Canli et al., 2002), and between Neuroticism and fearful stimuli (Cremers et al., 2010; Williams et al., 2006), there may be more-numerous null results for such associations which have not been published due to publication bias. On the other hand, the modality of the emotional stimulus may be important; for example, perhaps happy faces but not happy music elicits higher activation in subjects with higher Extraversion. Conversely, perhaps the separation of fearful and sad stimuli is important; some studies found that Neuroticism was related to brain activity during negative stimuli, but this category included sad and fearful stimuli (e.g., Canli et al., 2001).

### The regions-of-variance method

The purpose of the ROV method is to isolate regions of interest in which to investigate individual differences. Although the present study was not a systematic investigation of the effectiveness of this method, here we performed both ROV and whole-brain analyses in order to ascertain whether the ROV analysis was more sensitive and whether it would be consistent with robust whole-brain results. This was indeed demonstrated by both of the significant findings in this study. First, a similar cluster was found for Neuroticism and sad music in both the ROV and whole-brain analyses; it would be surprising if the ROV method had not found this result, since it was strong enough to be found in the more-conservative whole-brain analysis. However, it is notable that while the whole-brain cluster contained all of the voxels in the ROV cluster, the whole-brain cluster also included voxels not in the ROV cluster; indeed it was not possible for those voxels to be found in the ROV analysis because they were not included in the ROV map. This suggests that the threshold used on the ROV maps may not have been ideal; a more-lenient threshold may have included the entire whole-brain cluster in the ROV map. Future research on the ROV method could investigate how to threshold the maps in a way that optimizes the tradeoff between specificity and sensitivity. Second, the result for Openness and happy music suggests that the ROV method may be useful for improving the power of statistical tests to reveal brain areas that cannot be seen in the more-conservative whole-brain analysis; while the result for Openness and happy music did not pass the stringent cluster-size threshold for the whole-brain analysis, it did pass the threshold for the ROV analysis. Therefore, in this study, the ROV method was a useful tool for investigating individual differences, but there is room for improvement of this method.

## Conclusion

This study investigated the role of personality in neural activations to emotion in music. The results indicate a link between Neuroticism and neural activation in the inferior parietal lobe during perception of sad music; this trait-emotion conjunction is consistent with behavioral evidence that Neuroticism is related to negative-emotion tendencies (Larsen & Ketelaar, 1991), as well as previous brain research with non-musical stimuli suggesting that this trait is related to brain activity in limbic and multimodal processing areas during perception of negatively-valenced stimuli (Bruhl, Viebke, Baumgartner, Kaffenberger, & Herwig, 2011; Canli, Amin, Haas, Omura, & Constable, 2004; Canli et al., 2001; Cremers et al., 2010; Jimura, Konishi, & Miyashita, 2009; Omura et al., 2005). Further, this result points to the role of action observation network as a neural mechanism behind emotional contagion, mimicking and action tendencies for induction of emotional responses to music (Juslin & Västfjäll, 2008; Molnar-Szakacs & Overy, 2006). This result was found with both a regions-of-variance analysis as well as a whole-brain analysis. While no further results were found with the whole-brain analysis, the more-sensitive ROV analysis additionally found Openness to Experience to be related to neural activation in auditory and associative temporal regions in response to happy music. These results help elucidate the role of personality in emotion processing in the brain, and additionally, this study qualitatively demonstrates the sensitivity of the ROV method for investigating individual differences.

## References

Alluri, V., Toiviainen, P., Jääskeläinen, I. P., Glerean, E., Sams, M., & Brattico, E. (2012). Large-scale brain networks emerge from dynamic processing of musical timbre, key and rhythm. NeuroImage, 59 (4), 3677–3689.

Balkwill, L.-L., & Thompson, W. F. (1999). A cross-cultural investigation of the perception of emotion in music: Psychophysical and cultural cues. Music Perception: An Interdisciplinary Journal, 17 (1), 43–64.

Balkwill, L.-L., Thompson, W. F., & Matsunaga, R. (2004). Recognition of emotion in Japanese, Western, and Hindustani music by Japanese listeners. Japanese Psychological Research, 46 (4), 337–349.

Bogert, B., Numminen-Kontti, T., Gold, B., Sams, M., Numminen, J., Burunat, I., … Brattico, E. (2016). Hidden sources of joy, fear, and sadness: Explicit versus implicit neural processing of musical emotions. Neuropsychologia, 89, 393–402.

Bolger, N., & Schilling, E. A. (1991). Personality and the problems of everyday life: The role of Neuroticism in exposure and reactivity to daily stressors. Journal of Personality, 59 (3), 355–386.

Brattico, E., Alluri, V., Bogert, B., Jacobsen, T., Vartiainen, N., Nieminen, S. K., & Tervaniemi, M. (2011). A functional MRI study of happy and sad emotions in music with and without lyrics. Frontiers in Psychology, 2, 308.

Brattico, E., Bogert, B., Alluri, V., Tervaniemi, M., Eerola, T., & Jacobsen, T. (2016). It’s sad but I like it: The neural dissociation between musical emotions and liking in experts and laypersons. Frontiers in Human Neuroscience, 9, 676.

Brück, C., Kreifelts, B., Kaza, E., Lotze, M., & Wildgruber, D. (2011). Impact of personality on the cerebral processing of emotional prosody. Neuroimage, 58(1), 259–268.

Brühl, A. B., Viebke, M. C., Baumgartner, T., Kaffenberger, T., & Herwig, U. (2011). Neural correlates of personality dimensions and affective measures during the anticipation of emotional stimuli. Brain imaging and behavior, 5(2), 86–96.

Burger, B., Saarikallio, S., Luck, G., Thompson, M. R., & Toiviainen, P. (2013). Relationships between perceived emotions in music and music-induced movement. Music Perception: An Interdisciplinary Journal, 30 (5), 517–533.

Canli, T., Amin, Z., Haas, B., Omura, K., & Constable, R. T. (2004). A double dissociation between mood states and personality traits in the anterior cingulate. Behavioral Neuroscience, 118 (5), 897.

Canli, T., Sivers, H., Whitfield, S. L., Gotlib, I. H., & Gabrieli, J. D. (2002). Amygdala response to happy faces as a function of Extraversion. Science, 296 (5576), 2191–2191.

Canli, T., Zhao, Z., Desmond, J. E., Kang, E., Gross, J., & Gabrieli, J. D. (2001). An fMRI study of personality influences on brain reactivity to emotional stimuli. Behavioral Neuroscience, 115 (1), 33.

Carlson, E., Saarikallio, S., Toiviainen, P., Bogert, B., Kliuchko, M., & Brattico, E. (2015). Maladaptive and adaptive emotion regulation through music: A behavioral and neuroimaging study of males and females. Frontiers in Human Neuroscience, 9, 466.

Caspers, S., Zilles, K., Laird, A. R., & Eickhoff, S. B. (2010). ALE meta-analysis of action observation and imitation in the human brain. NeuroImage, 50 (3), 1148–1167.

Cloninger, C. R. (2000). Biology of personality dimensions. Current Opinion in Psychiatry, 13 (6), 611–616.

Coan, J. A., & Allen, J. J. (2003). Frontal EEG asymmetry and the behavioral activation and inhibition systems. Psychophysiology, 40 (1), 106–114.

Colver, M. C., & El-Alayli, A. (2016). Getting aesthetic chills from music: The connection between Openness to Experience and frisson. Psychology of Music, 44 (3), 413–427.

Concina, G., Renna, A., Grosso, A., & Sacchetti, B. (2019). The auditory cortex and the emotional valence of sounds. Neuroscience & Biobehavioral Reviews.

Costa, P. T., & McCrae, R. R. (1980). Influence of Extraversion and Neuroticism on subjective well-being: Happy and unhappy people. Journal of Personality and Social Psychology, 38 (4), 668.

Costa, P. T., & McCrea, R. R. (1992). Revised NEO Personality Inventory (NEO PI-R) and NEO Five-Factor Inventory (NEO-FFI). Psychological Assessment Resources.

Costa Jr, P. T., & McCrae, R. R. (1992). NEO personality inventory–revised (NRO PI-R) and NEO five-factor inventory (NEO-FFI) professional manual. Odessa, FL: Psychological Assessment Resources.

Craig, A., Tran, Y., Hermens, G., Williams, L. M., Kemp, A., Morris, C., & Gordon, E. (2009). Psychological and neural correlates of emotional intelligence in a large sample of adult males and females. Personality and Individual Differences, 46 (2), 111–115.

Cremers, H. R., Demenescu, L. R., Aleman, A., Renken, R., van Tol, M. J., van der Wee, N. J., … Roelofs, K. (2010). Neuroticism modulates amygdala-prefrontal connectivity in response to negative emotional facial expressions. NeuroImage, 49 (1), 963–970.

David, J. P., Green, P. J., Martin, R., & Suls, J. (1997). Differential roles of Neuroticism, Extraversion, and event desirability for mood in daily life: An integrative model of top-down and bottom-up influences. Journal of Personality and Social Psychology, 73 (1), 149.

Deng, Y., Li, S., Zhou, R., & Walter, M. (2018). Motivation but not valence modulates neuroticism-dependent cingulate cortex and insula activity. Human brain mapping, 39(4), 1664–1672.

Eerola, T., & Vuoskoski, J. K. (2011). A comparison of the discrete and dimensional models of emotion in music. Psychology of Music, 39 (1), 18–49.

Eklund, A., Nichols, T. E., & Knutsson, H. (2016). Cluster failure: Why fMRI inferences for spatial extent have inflated false-positive rates. Proceedings of the National Academy of Sciences, 113 (28), 7900–7905.

Ethofer, T., Anders, S., Erb, M., Herbert, C., Wiethoff, S., Kissler, J., … Wildgruber, D. (2006). Cerebral pathways in processing of affective prosody: A dynamic causal modeling study. NeuroImage, 30 (2), 580–587.

Eysenck, H. J. (1967). The biological basis of personality. Thomas.

Fisher, R. A. (1915). Frequency distribution of the values of the correlation coefficient in samples from an indefinitely large population. Biometrika, 10 (4), 507–521.

Fritz, T., Jentschke, S., Gosselin, N., Sammler, D., Peretz, I., Turner, R., … Koelsch, S. (2009). Universal recognition of three basic emotions in music. Current Biology, 19 (7), 573–576.

Gignac, G. E., & Szodorai, E. T. (2016). Effect size guidelines for individual differences researchers. Personality and Individual Differences, 102, 74–78.

Goldin, P. R., McRae, K., Ramel, W., & Gross, J. J. (2008). The neural bases of emotion regulation: Reappraisal and suppression of negative emotion. Biological Psychiatry, 63 (6), 577–586.

Gray, J. A. (1984). The neuropsychology of anxiety. In Fortschritte der experimentalpsychologie (pp. 52–71). Springer.

Gross, J. J., & John, O. P. (2003). Individual differences in two emotion regulation processes: Implications for affect, relationships, and well-being. Journal of Personality and Social Psychology, 85 (2), 348.

Haas, B. W., Constable, R. T., & Canli, T. (2008). Stop the sadness: Neuroticism is associated with sustained medial prefrontal cortex response to emotional facial expressions. NeuroImage, 42 (1), 385–392.

Harmon-Jones, E., & Allen, J. J. B. (1997). Behavioral activation sensitivity and resting frontal EEG asymmetry: Covariation of putative indicators related to risk for mood disorders. Journal of Abnormal Psychology, 106 (1), 159.

Holmes, A., & Friston, K. (1998). Generalisability, random effects & population inference. NeuroImage, 7, 754.

Hutcherson, C. A., Goldin, P. R., Ramel, W., McRae, K., & Gross, J. J. (2008). Attention and emotion influence the relationship between extraversion and neural response. Social Cognitive and Affective Neuroscience, 3(1), 71–79.

Jacoby, N., & McDermott, J. H. (2017). Integer ratio priors on musical rhythm revealed cross-culturally by iterated reproduction. Current Biology, 27 (3), 359–370.

Jimura, K., Konishi, S., & Miyashita, Y. (2009). Temporal pole activity during perception of sad faces, but not happy faces, correlates with Neuroticism trait. Neuroscience Letters, 453 (1), 45–48.

John, O. P., & Srivastava, S. (1999). The Big Five trait taxonomy: History, measurement, and theoretical perspectives. In Handbook of personality: Theory and research (Vol. 2, pp. 102–138). Guilford.

Jorm, A. F., Christensen, H., Henderson, A. S., Jacomb, P. A., Korten, A. E., & Rodgers, B. (1998). Using the BIS/BAS scales to measure behavioural inhibition and behavioural activation: Factor structure, validity and norms in a large community sample. Personality and Individual Differences, 26 (1), 49–58.

Juslin, P. N., & Laukka, P. (2004). Expression, perception, and induction of musical emotions: A review and a questionnaire study of everyday listening. Journal of New Music Research, 33 (3), 217–238.

Juslin, P. N., & Västfjäll, D. (2008). Emotional responses to music: The need to consider underlying mechanisms. Behavioral and Brain Sciences, 31 (5), 559–575.

Jylhä, P., & Isometsä, E. (2006). The relationship of Neuroticism and Extraversion to symptoms of anxiety and depression in the general population. Depression and Anxiety, 23 (5), 281–289.

Kehoe, E. G., Toomey, J. M., Balsters, J. H., & Bokde, A. L. (2011). Personality modulates the effects of emotional arousal and valence on brain activation. Social Cognitive and Affective Neuroscience, 7(7), 858–870.

Klamer, S., Schwarz, L., Krüger, O., Koch, K., Erb, M., Scheffler, K., & Ethofer, T. (2017). Association between neuroticism and emotional face processing. Scientific reports, 7(1), 17669.

Koelsch, S. (2014). Brain correlates of music-evoked emotions. Nature Reviews Neuroscience, 15 (3), 170.

Koelsch, S. (2018). Investigating the neural encoding of emotion with music. Neuron, 98 (6), 1075–1079.

Koelsch, S., Fritz, T., v Cramon, D. Y., Müller, K., & Friederici, A. D. (2006). Investigating emotion with music: An fMRI study. Human Brain Mapping, 27 (3), 239–250.

Koelsch, S., Skouras, S., & Jentschke, S. (2013). Neural correlates of emotional personality: A structural and functional magnetic resonance imaging study. PLoS One, 8 (11), e77196.

Koelsch, S., Skouras, S., & Lohmann, G. (2018). The auditory cortex hosts network nodes influential for emotion processing: An fmri study on music-evoked fear and joy. PloS one, 13 (1), e0190057.

Kotz, S. A., Meyer, M., Alter, K., Besson, M., von Cramon, D. Y., & Friederici, A. D. (2003). On the lateralization of emotional prosody: An event-related functional MR investigation. Brain and Language, 86 (3), 366–376.

Ladinig, O., & Schellenberg, E. G. (2012). Liking unfamiliar music: Effects of felt emotion and individual differences. Psychology of Aesthetics, Creativity, and the Arts, 6 (2), 146.

Larsen, R. J., & Ketelaar, T. (1991). Personality and susceptibility to positive and negative emotional states. Journal of Personality and Social Psychology, 61 (1), 132.

Lundqvist, L.-O., Carlsson, F., Hilmersson, P., & Juslin, P. N. (2009). Emotional responses to music: Experience, expression, and physiology. Psychology of Music, 37 (1), 61–90.

Marco, C. A., & Suls, J. (1993). Daily stress and the trajectory of mood: Spillover, response assimilation, contrast, and chronic negative affectivity. Journal of Personality and Social Psychology, 64 (6), 1053.

McDermott, J. H., Schultz, A. F., Undurraga, E. A., & Godoy, R. A. (2016). Indifference to dissonance in native Amazonians reveals cultural variation in music perception. Nature, 535 (7613), 547.

Meister, I. G., Krings, T., Foltys, H., Boroojerdi, B., Müller, M., Töpper, R., & Thron, A. (2004). Playing piano in the mind - An fMRI study on music imagery and performance in pianists. Cognitive Brain Research, 19 (3), 219–228.

Mitterschiffthaler, M. T., Fu, C. H., Dalton, J. A., Andrew, C. M., & Williams, S. C. (2007). A functional MRI study of happy and sad affective states induced by classical music. Human Brain Mapping, 28 (11), 1150–1162.

Molnar-Szakacs, I., & Overy, K. (2006). Music and mirror neurons: From motion to ‘emotion. Social cognitive and affective neuroscience, 1 (3), 235–241.

Nichols, T. E., & Holmes, A. P. (2002). Nonparametric permutation tests for functional neuroimaging: A primer with examples. Human Brain Mapping, 15 (1), 1–25.

North, A. C., Hargreaves, D. J., & Hargreaves, J. J. (2004). Uses of music in everyday life. Music Perception: An Interdisciplinary Journal, 22 (1), 41–77.

Olson, I. R., Plotzker, A., & Ezzyat, Y. (2007). The enigmatic temporal pole: A review of findings on social and emotional processing. Brain, 130 (7), 1718–1731.

Omura, K., Aron, A., & Canli, T. (2005). Variance maps as a novel tool for localizing regions of interest in imaging studies of individual differences. Cognitive, Affective, & Behavioral Neuroscience, 5 (2), 252–261.

Park, M., Hennig-Fast, K., Bao, Y., Carl, P., Pöppel, E., Welker, L., … Gutyrchik, E. (2013). Personality traits modulate neural responses to emotions expressed in music. Brain Research, 1523, 68–76.

Pavot, W., Diener, E., & Fujita, F. (1990). Extraversion and happiness. Personality and Individual Differences, 11 (12), 1299–1306.

Poldrack, R. A. (2007). Region of interest analysis for fMRI. Social cognitive and affective neuroscience, 2 (1), 67–70.

Rentfrow, P. J., & Gosling, S. D. (2003). The do re mi’s of everyday life: The structure and personality correlates of music preferences. Journal of Personality and Social Psychology, 84 (6), 1236.

Richard, E., & Diener, E. (2009). Personality and subjective well-being. In E. Diener (Ed.), The science of well-being (pp. 75–102). Springer.

Samson, D., Apperly, I. A., Chiavarino, C., & Humphreys, G. W. (2004). Left temporoparietal junction is necessary for representing someone else’s belief. Nature Neuroscience, 7 (5), 499.

Sander, D., Grandjean, D., Pourtois, G., Schwartz, S., Seghier, M. L., Scherer, K. R., & Vuilleumier, P. (2005). Emotion and attention interactions in social cognition: Brain regions involved in processing anger prosody. NeuroImage, 28 (4), 848–858.

Saxe, R., & Kanwisher, N. (2003). People thinking about thinking people: The role of the temporo-parietal junction in “theory of mind”. NeuroImage, 19 (4), 1835–1842.

Scherer, K. R. (2005). What are emotions? And how can they be measured? Social Science Information, 44 (4), 695–729.

Schmithorst, V. J., & Holland, S. K. (2003). The effect of musical training on music processing: A functional magnetic resonance imaging study in humans. Neuroscience Letters, 348 (2), 65–68.

Schnell, K., Bluschke, S., Konradt, B., & Walter, H. (2011). Functional relations of empathy and mentalizing: An fMRI study on the neural basis of cognitive empathy. NeuroImage, 54 (2), 1743–1754.

Servaas, M. N., Riese, H., Ormel, J., & Aleman, A. (2014). The neural correlates of worry in association with individual differences in Neuroticism. Human Brain Mapping, 35 (9), 4303–4315.

Silvia, P. J., Fayn, K., Nusbaum, E. C., & Beaty, R. E. (2015). Openness to Experience and awe in response to nature and music: Personality and profound aesthetic experiences. Psychology of Aesthetics, Creativity, and the Arts, 9 (4), 376.

Sloboda, J. A., O’Neill, S. A., & Ivaldi, A. (2001). Functions of music in everyday life: An exploratory study using the experience sampling method. Musicae Scientiae, 5 (1), 9–32.

Suslow, T., Kugel, H., Reber, H., Bauer, J., Dannlowski, U., Kersting, A., … & Egloff, B. (2010). Automatic brain response to facial emotion as a function of implicitly and explicitly measured extraversion. Neuroscience, 167(1), 111–123.

Vallar, G., & Papagno, C. (1995). Neuropsychological impairments of short-term memory. John Wiley & Sons.

Vuoskoski, J. K., & Eerola, T. (2011a). Measuring music-induced emotion: A comparison of emotion models, personality biases, and intensity of experiences. Musicae Scientiae, 15 (2), 159–173.

Vuoskoski, J. K., & Eerola, T. (2011b). The role of mood and personality in the perception of emotions represented by music. Cortex, 47 (9), 1099–1106.

Vuoskoski, J. K., Thompson, W. F., McIlwain, D., & Eerola, T. (2012). Who enjoys listening to sad music and why? Music Perception: An Interdisciplinary Journal, 29 (3), 311–317.

Wang, L., Shi, Z., & Li, H. (2009). Neuroticism, Extraversion, emotion regulation, negative affect and positive affect: The mediating roles of reappraisal and suppression. Social Behavior and Personality: An International Journal, 37 (2), 193–194.

Williams, L. M., Brown, K. J., Palmer, D., Liddell, B. J., Kemp, A. H., Olivieri, G., … Gordon, E. (2006). The mellow years?: Neural basis of improving emotional stability over age. Journal of Neuroscience, 26 (24), 6422–6430.

Zatorre, R. J., Evans, A. C., Meyer, E., & Gjedde, A. (1992). Lateralization of phonetic and pitch discrimination in speech processing. Science, 256 (5058), 846–849.

