## Supplementary Materials; Figure S1; Table S1 for "Personality modulates brain responses to emotion in music: Comparing whole-brain and regions-of-variance approaches"

For the following paper:

#### Literature search

A literature search was conducted in order to review how previous studies investigated the role of personality in activations to emotional stimuli and what were their results. Three systematic searches were run, and additional material was found by looking in references and unsystematic searching.

The first systematic search was performed using Ovid. Specifically, the database PsycINFO <1987 to January Week 4 2019> was queried using keywords with the following search strategy:

1. exp EXTRAVERSION/ (2664)
2. neuroticism/ (3824)
3. openness to experience/ (922) 4. 1 or 2 or 3 (5930)
4. emotions/ or exp emotional states/ or exp emotional style/ or exp negative emotions/ or exp positive emotions/ (250312)
5. affective valence/ (2315) 7. 5 or 6 (251790)
6. 4 and 7 (1293)
7. exp brain/ (217792)
8. electrical activity/ (11432)
9. functional magnetic resonance imaging/ (19932)
10. magnetic resonance imaging/ (18631) 13. 9 or 11 or 12 (234910)
11. 8 and 13 (38)

The second systematic search was also a keyword search using MeSH terms on PubMed, using the following syntax: *((((”Neuroimaging”[Mesh]) OR ”Magnetic Resonance Imaging”[Mesh]) OR ”Brain”[Mesh])) AND (((((”openness to experience”) OR ”Neuroticism”[Mesh]) OR ”Extraversion (Psychology)”[Mesh])) AND ”Emotions”[Mesh]).* Note that there was no MeSH category for Openness to Experience, and so it was searched as a regular text word.

A third systematic search was done on PubMed. The purpose of this second search was to look through titles, abstracts, and author-provided keywords in recent publications that may not have been assigned database keywords or MeSH categories yet. The search was limited to the publication dates between 2016/01/01 and 2019/01/29, and the following syntax was used: *((((((extraver*[Text Word]) OR extrover*[Text Word]) OR neurotic*[Text Word]) OR openness[Text Word])) AND (((((brain[Text Word]) OR neuroimag*[Text Word]) OR fmri[Text Word]) OR ”functional magnetic resonance imag* *ing"[Text Word]) OR "functional mri"[Text Word])) AND ((emotion*[Text Word]) OR affect*[Text Word]).*

The studies found in these searches had to meet all of following inclusion criteria in order to be included in the review:

- Investigates brain activation using functional magnetic resonance imaging,
- Investigates explicitly-measured Neuroticism, Extraversion, and/or Openness to Experience, and
- Uses contrasting affective/emotional stimuli.

Fifteen studies met these criteria. All but one are listed in Tables 1 and 2 in the main document; the study by Suslow, Kugel, Lindner, Dannlowski, and Egloff (2017) met the inclusion criteria, but it had no significant results for explicitly-measured extraversion when trait anxiety was controlled for, and so it was not included in the table.

#### Analyses

##### Creation of stimulus regressors

For each subject and condition, a boxcar function was created describing when music of the given stimulus type was heard by that participant; this function was set to 1 when the given stimulus type was presented, and 0 elsewhere. These boxcar functions were then up-sampled by 10 (giving 20 time bins per scan) and were convolved with a double-gamma hemodynamic response function that was likewise up-sampled. The resulting convolved stimulus functions were then down-sampled to match the data sampling rate.

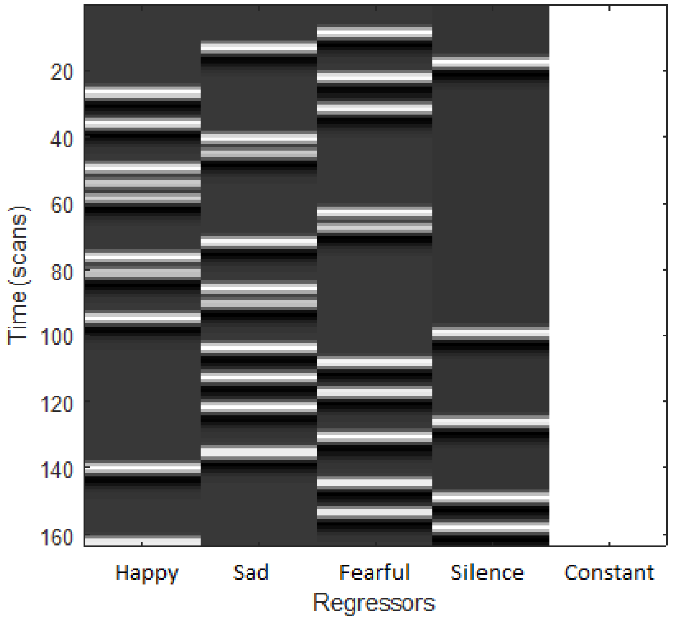


Figure S1: Graphical illustration of one participant's level-1 design matrix

The same method was used to create a function that described when 5 periods of silence occurred for each participant; these periods were not of interest to the current study, but they were included in the model as a regressor of no interest. The answer periods were not explicitly modeled to avoid over-parametrization. See Figure S1 for an example design matrix for one participant, which consists of three stimulus regressors, a silence regressor, and a constant.

##### Contrast calculations

With the parameter weights in the order of Happy, Sad, Fearful, Silence, and Constant (column of ones), the contrast vectors for each condition were as follows:

[2 -1 -1 0 0] for the Happy condition,
[-1 2 -1 0 0] for the Sad condition, and
[-1 -1 2 0 0] for the Fearful condition.

##### Calculation of variance maps

The equations for calculating the variance maps from which ROVs are selected are as follows (Omura, Aron, & Canli, 2005):

$$\boldsymbol{S_B^2} = \frac{\frac{\mathbf{1}}{\boldsymbol{Nsubj-1}} \sum_{i=1}^{Nsubj} (\boldsymbol{con*.img_i} - \overline{\boldsymbol{con*.img}})}{\mathbf{2}} * \boldsymbol{Nscan} \quad (1)$$

$$\boldsymbol{S_W^2} = \frac{\frac{\mathbf{1}}{\boldsymbol{Nsubj}} \sum_{i=1}^{Nsubj} (\boldsymbol{ResMS.img_i})}{\boldsymbol{Nscan-1}} \quad (2)$$

$$\boldsymbol{F} = \frac{\boldsymbol{S_W^2}}{\boldsymbol{S_B^2}} \quad (3)$$

Where

$S_B^2$ is the between-subjects variance,

$S_W^2$ is the within-subjects variance,

$F$ is the ratio of between- to within-subjects variance,

$Nsubj$ is the total number of subjects,

$Nscan$ is the total number of scans,

$con*.img_i$ is the contrast image for the given participant,

$\overline{con*.img}$ is the mean contrast image (mean across participants), and

$ResMS.img_i$ is the image of the mean squared residual (mean across time) for the given participant.

Essentially, for each voxel, an *F* -value was calculated, which is the ratio of the between-subjects variance in the level-1 parameter estimates to the mean within-subjects residual variance for that voxel (mean across participants). These *F* -values make up the variance maps.

**Thresholding variance maps to isolate regions of variance (ROVs)**

For each emotion condition, voxels in the *F* -map were sorted in a vector according to their *F* -value, and then the 'change point' in the vector was determined using the *findchangepts* algorithm in MATLAB, with the *linear* statistic option. *Findchangepts* uses a parametric global method to find abrupt changes in a curve. The change point is the point at which the total residual error in the chosen statistic is at a minimum, that is, the sum of a) the deviation of the points prior to the change point from the summary statistic over those prior points (e.g., the mean) and b) the deviation of the points subsequent to the change point from the summary statistic over those subsequent points. The *linear* statistic option uses a summary statistic describing both the mean and the slope of the line. The sorted *F* -voxel-values for each condition and the respective change point for each line is shown in Figure S2.

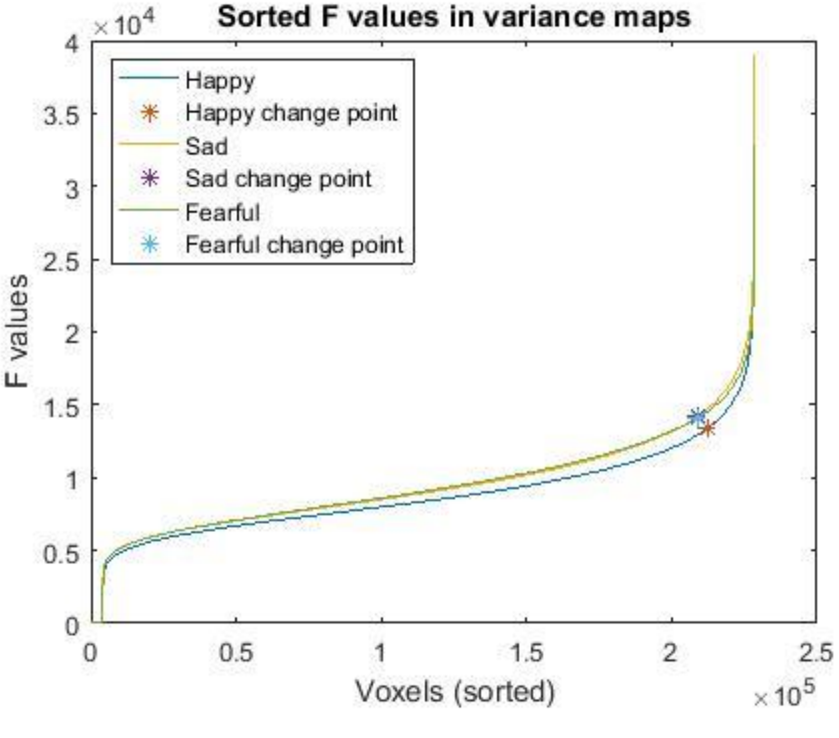


Figure S2: Sorted *F* -values in variance maps for each condition, and change point in each line (*), which is used as the cluster-forming threshold for the respective condition's variance map.

Thus, the ROV maps were height-thresholded at the voxel level using this change-point as the threshold. The maps were then subjected to an arbitrary cluster-size threshold of 15 voxels and were binarized in order to be used as masks the first-level contrast betas. The ROV map for each condition can be seen in Figure S3.

As far as we are aware, no previous studies have thresholded statistical parametric maps using this method of sorting statistics and finding the change in the line. It bears some resemblance to the scree-plot method used for selecting partitions in dimensionality-reduction methods such as factor analysis and principal component analysis. It may seem unusual that there is no significance level attached to these thresholds. However, the choice of height and extent thresholds for the ROV maps was not considered to be of great importance because a data-driven threshold was used for significance estimation of the level-2 model; the conservativeness of the level-2 significance threshold would be inversely related to the conservativeness of these ROV thresholds. For example, using *more*-conservative thresholds for creating the ROV masks would result in maps with fewer unmasked voxels; this means that the level-2 analysis would be performed on fewer voxels and there would be fewer comparisons to correct for when correcting for multiple comparisons, resulting in a *less*-conservative level-2 significance test. That being said, it is not known whether this relationship is linear, so there may in fact be ideal cluster- height and -extent thresholds for the ROV maps that optimally balance Type I and Type II errors in the level-2 results. This was not investigated in the present study.

**Avoiding circular analysis and 'double dipping' in functional ROI selection**

Methods of functionally deriving ROIs run the risk of circularity; significance may be overestimated due to 'double dipping' when the same data is used to define regions of interest for an analysis and to perform the analysis (Kriegeskorte, Simmons, Bellgowan, & Baker, 2009). Such methods increase false positive rates by preferentially selecting noise that may be in line with the desired effect. This criticism has been made for the method of selecting ROIs functionally based on common activation across participants: if you are looking for consistent neural activity across subjects, then selecting regions that are consistently activated across subjects will make it more likely that you find the hypothesized result by chance, as you are preferentially selecting noise that is more likely to be in line with any hypothesis of a significant mean activation.

The relevant question here is whether functionally deriving ROIs as regions of *variance* runs the same risk of circularity. Here I argue that this is not the case. Selecting regions with similar activation between participants increases the likelihood of the mean activation being significant by chance, hence the 'double dipping'. But in the context of the ROV method, selecting regions with high between-subjects variance (relative to within-subjects variance) could select noise that is by chance related to the behavioral individual difference of interest, however, it could also select noise that is highly variable between participants but is unrelated to the individual difference. One could even argue that there are more ways for noise to be randomly *un*related to a given individual difference than to be randomly related.

Therefore, functionally selecting regions of variance should not necessarily increase the false-positive rate. Perhaps future research could demonstrate this more convincingly using proofs and simulations. But in the present study, extra precaution was taken to avoid double-dipping by using a nonparametric method of significance estimation. This method involves building a distribution of null results using the data within the regions of variance, and so it controls for any potential increase in false positives resulting from the ROI-selection procedure.

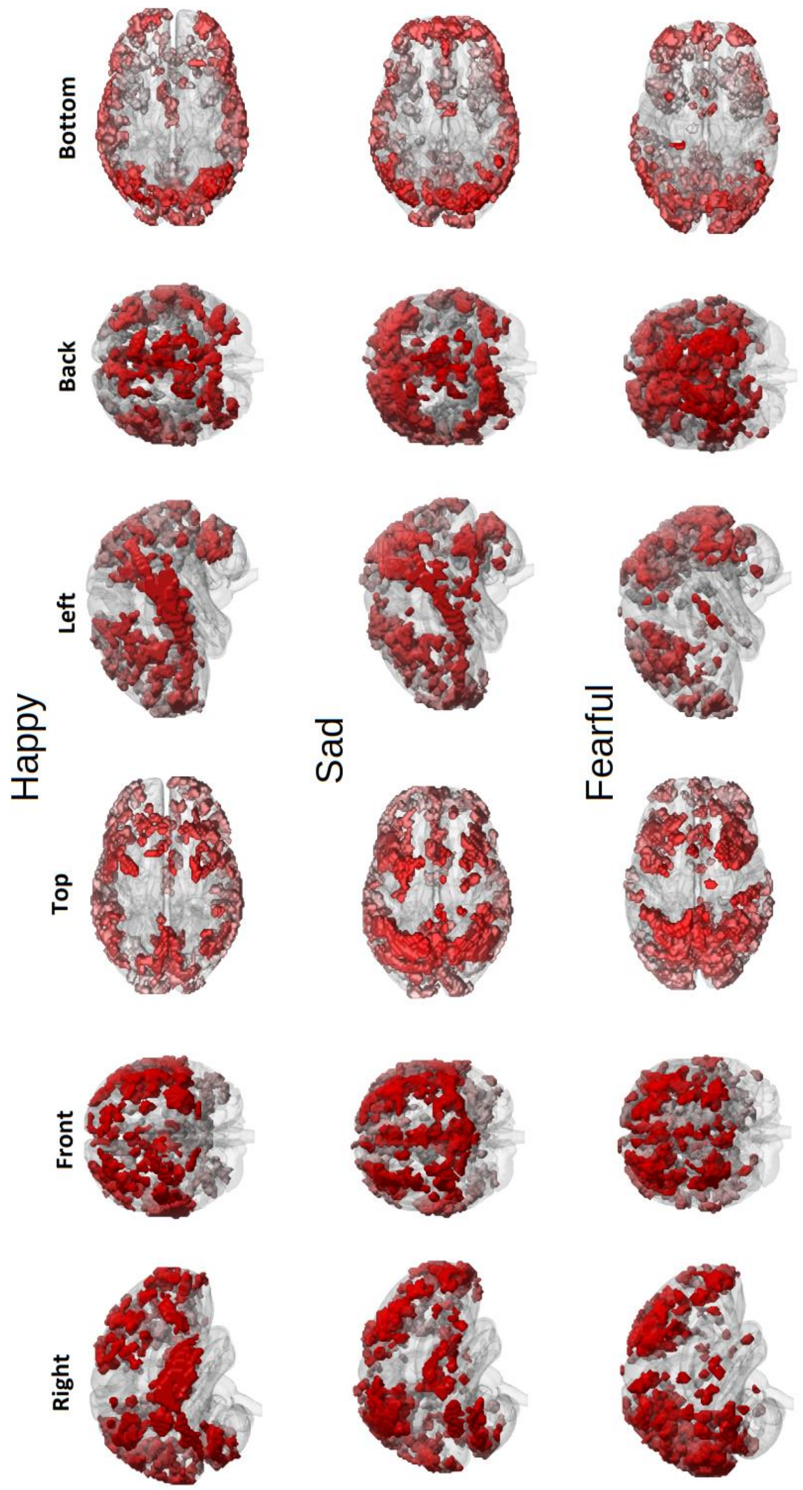


Figure S3: Regions-of-Variance maps for each of the emotional music conditions.

**Statistical inference and correction for multiple comparisons**

A Monte Carlo simulation was carried out on the Z-maps, in order to find a cluster-level threshold that corrected for multiple comparisons at a FWER of 5%. In order to control for FWE in a nonparametric manner, one needs to build a null distribution of maximum values of the statistic being used (e.g., voxel *p*-values or cluster sizes/masses), and then select the $n^{th}$ statistic in the distribution as the threshold, $n$ being $\lfloor (1 - FWER) * NumberOfPermutations \rfloor$ ($\lfloor\ \rfloor$ means rounded down). In other words, if you want to control for FWER at 5%, and you have run 10000 permutations, then the cluster-statistic threshold is the $9500^{th}$ number in the ascending-sorted list of statistics.

The reason why one stores only the maximum statistic from each permutation, and not all the statistics, is that FWE occurs when at least *one* voxel/cluster statistic is a false positive. If two statistics are over the FWER threshold, those two statistics will be the maximum and next-to-maximum statistics. If one statistic is over the threshold, it will be the maximum statistic. If no statistics are over the threshold, then the maximum is not over the threshold. Thus, the FWER threshold is derived from a distribution of maximum statistics.

Nichols and Holmes (2002) outlined the steps for performing a permutation test for significance testing in neuroimaging. Here are the steps and how they were carried out in the present study:

1. *Null Hypothesis:* The null hypothesis is that there is no correlation between the given personality trait and the set of contrast betas from the given condition.

2. *Exchangeability:* Under the null hypothesis, participants are exchangeable; if there is no correlation, then exchanging individual participants' data should not make a difference to the results. Therefore, participants' data can be randomized in order to build a null distribution of cluster statistics.

3. *Statistic:* Permutations were run for three different voxel-level cluster-forming thresholds: $p < .05$, .01, and .001. The reason for using multiple thresholds is that there is not a clear answer as to which threshold is ideal. Two types of summary statistic were calculated for each cluster: a) cluster size, which is the number of voxels in the cluster, and b) cluster mass, which is the sum of the Z-values in the cluster. The reason for calculating both types of summary statistics is that cluster size is a more conventional measure, and cluster mass may be more intuitive because it can equates a lowly-activated large cluster with a highly-activated small cluster.

4. *Relabeling:* Ideally, one would do a permutation for every possible relabeling of the participants, but with 55 participants the $1.27 \times 10^{73}$ possible permutations is not computationally realistic. When there are too many possible permutations, a subset can used (Dwass, 1957; Edgington, 1969), in what is known as an approximate test, Monte-Carlo permutation test, or random permutation test. In this case, 10000 permutations were used.

5. *Permutation Distribution:* For each of the 10000 permutations, the sets of 3 trait scores for each participant were randomized across participants (i.e., participant labels were randomized). In order to ensure that each randomization was unique, the *rng('shuffle')* MATLAB command was called before the parallelized permutation loop. For each combination of personality trait and emotional condition, the level-2 partial correlation was calculated between the given trait and the contrast betas for the given condition, controlling for the remaining traits. The correlation coefficients were normalized using the Fisher transform. For the regions-of-variance analysis, the Z-maps were masked with the ROV map corresponding to the given condition (previously calculated with the real level-1 data). No masking was applied for the whole-brain analysis. For each height threshold ($p$ < .05, .01, and .001), the maps were thresholded with a voxelwise cluster-forming threshold at the corresponding 2-tailed Z-value (i.e., Z = 1.96, 2.58, and 3.29). For two voxels to be in the same cluster, they needed to have one side, edge, or corner touching. For every cluster passing the threshold, the cluster size and mass was calculated, and then the maximum size and mass across clusters was saved. In this way, a null distribution of maximum statistics was built for every combination of trait, condition, and height threshold, for both the ROV and whole-brain analyses and for both the cluster-size and -mass statistics (i.e., 3 x 3 x 3 x 2 x 2 = 108 distributions).

6. *Significance:* For each distribution, the 10000 values were sorted in ascending order. Then the cluster-statistic threshold was selected as the 9500*th* number, corresponding to a FWER of 5%. Each threshold was then applied to the corresponding map of the real data.

See Table S1 for the cluster-size and -mass thresholds that were derived in this way.

Table S1: Cluster-size (number of voxels) and -mass (sum of Z-scores) thresholds for each combination of personality trait, condition, and significance threshold, and for both the region-of-variance and whole-brain analyses. CLF: Cluster-Forming Threshold. ROV: Regions of Variance.

| CLUSTER-FORMING THRESHOLD | | CLUSTER SIZE | | CLUSTER MASS | |
|---|---|---|---|---|---|
| | | ROV | Whole-brain | ROV | Whole-brain |
| **EXTRAVERSION** | | | | | |
| *Happy > Sad & Fear* | | | | | |
| | *p* < .050 | 491 | 15 126 | 251.60 | 37 050.27 |
| | *p* < .010 | 170 | 1 383 | 512.55 | 4 158.78 |
| | *p* < .001 | 42 | 180 | 152.07 | 665.44 |
| *Sad > Happy & Fear* | | | | | |
| | *p* < .050 | 801 | 14 746 | 1 992.42 | 36 329.86 |
| | *p* < .010 | 254 | 1 446 | 764.35 | 4 388.87 |
| | *p* < .001 | 50 | 183 | 182.97 | 678.52 |
| *Fear > Happy & Sad* | | | | | |
| | *p* < .050 | 1 080 | 19 407 | 2 718.17 | 47 342.37 |
| | *p* < .010 | 263 | 1 560 | 782.34 | 4 696.45 |
| | *p* < .001 | 46 | 176 | 167.57 | 650.13 |
| **NEUROTICISM** | | | | | |
| *Happy > Sad & Fear* | | | | | |
| | *p* < .050 | 503 | 14 530 | 1 302.21 | 35 471.82 |
| | *p* < .010 | 189 | 1 433 | 578.84 | 4 402.41 |
| | *p* < .001 | 54 | 209 | 196.69 | 777.25 |
| *Sad > Happy & Fear* | | | | | |
| | *p* < .050 | 818 | 13 644 | 2 042.87 | 33 588.34 |
| | *p* < .010 | 272 | 1 487 | 837.22 | 4 510.42 |
| | *p* < .001 | 68 | 207 | 248.29 | 760.68 |
| *Fear* | | | | | |
| | *p* < .050 | 987 | 19 203 | 2 472.52 | 46 930.59 |
| | *p* < .010 | 254 | 1 621 | 773.39 | 4 913.25 |
| | *p* < .001 | 59 | 201 | 216.90 | 741.21 |
| **OPENNESS** | | | | | |
| *Happy > Sad & Fear* | | | | | |
| | *p* < .050 | 507 | 15 137 | 1 295.28 | 37 107.89 |
| | *p* < .010 | 182 | 1 445 | 559.85 | 4 392.54 |
| | *p* < .001 | 49 | 194 | 179.83 | 724.62 |
| *Sad > Happy & Fear* | | | | | |
| | *p* < .050 | 781 | 14 352 | 1990.24 | 35 319.33 |
| | *p* < .010 | 253 | 1 484 | 775.25 | 4 478.00 |
| | *p* < .001 | 59 | 193 | 216.30 | 709.37 |
| *Fear > Happy & Sad* | | | | | |
| | *p* < .050 | 1 018 | 19 825 | 2 528.13 | 48 778.74 |
| | *p* < .010 | 260 | 1 743 | 787.52 | 5 222.47 |
| | *p* < .001 | 54 | 197 | 197.57 | 5 222.47 |

#### Results

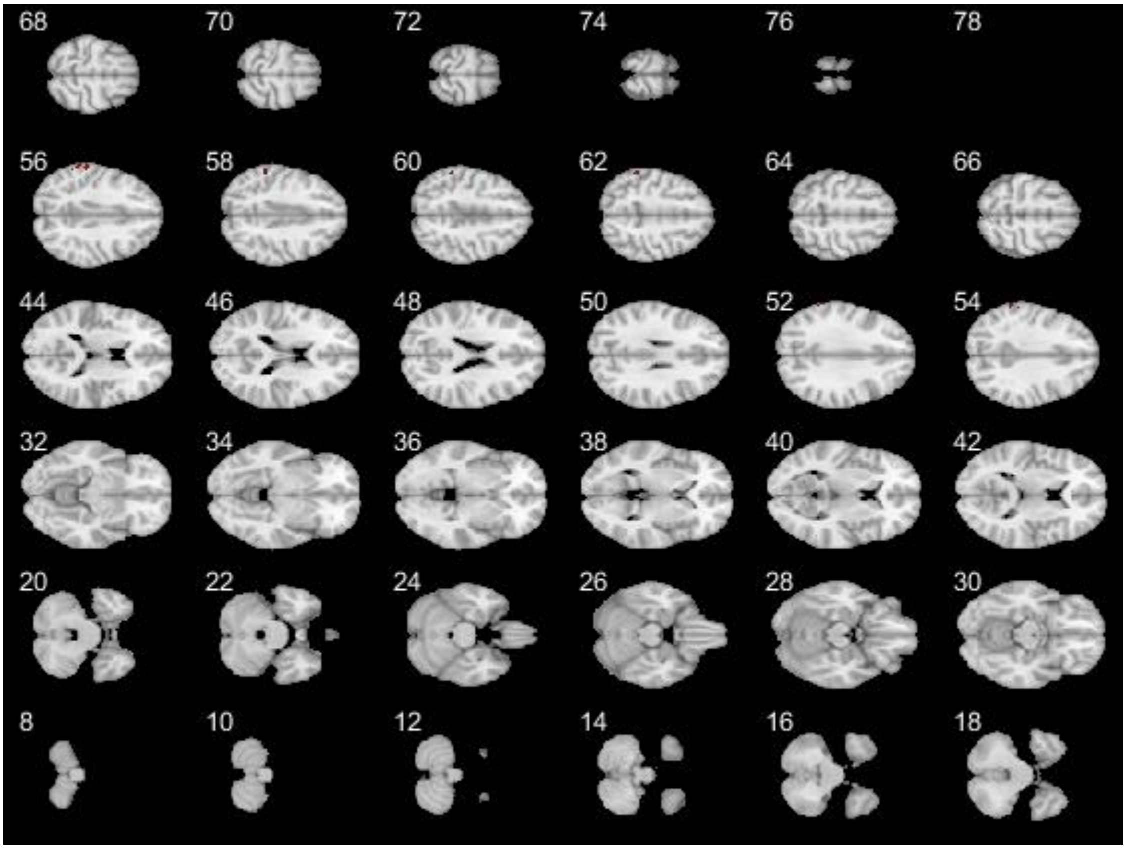


Figure S4: Significant cluster for Neuroticism & Sad music using the ROV analysis. This result was found with a CDT of $p = 0.001$, corrected at 5% FWER using non-parametric cluster-size and -mass thresholds, which were $k = 68$ voxels and Z = 248.49, respectively.

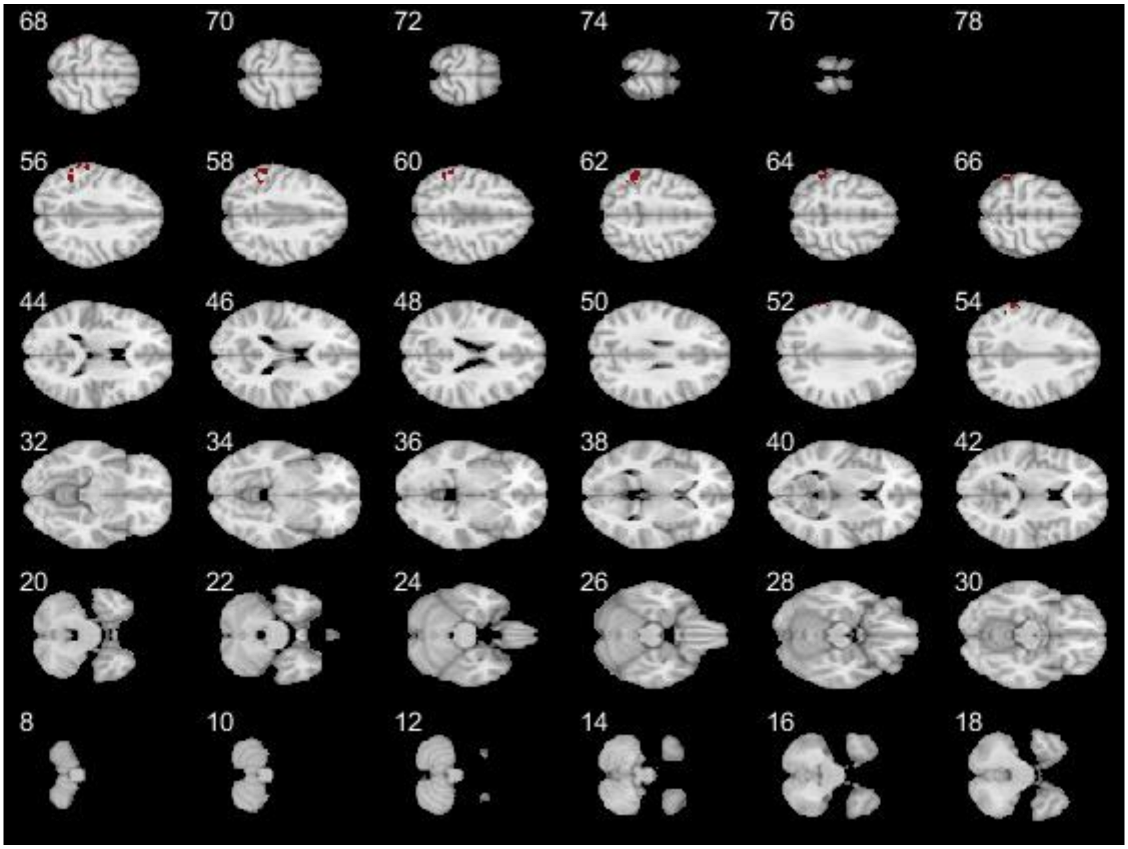


Figure S5: Significant cluster for Neuroticism & Sad music using the whole-brain analysis. This result was found with a CDT of $p = 0.001$, corrected at 5% FWER using non-parametric cluster-size and -mass thresholds, which were $k = 207$ voxels and Z = 760.68.

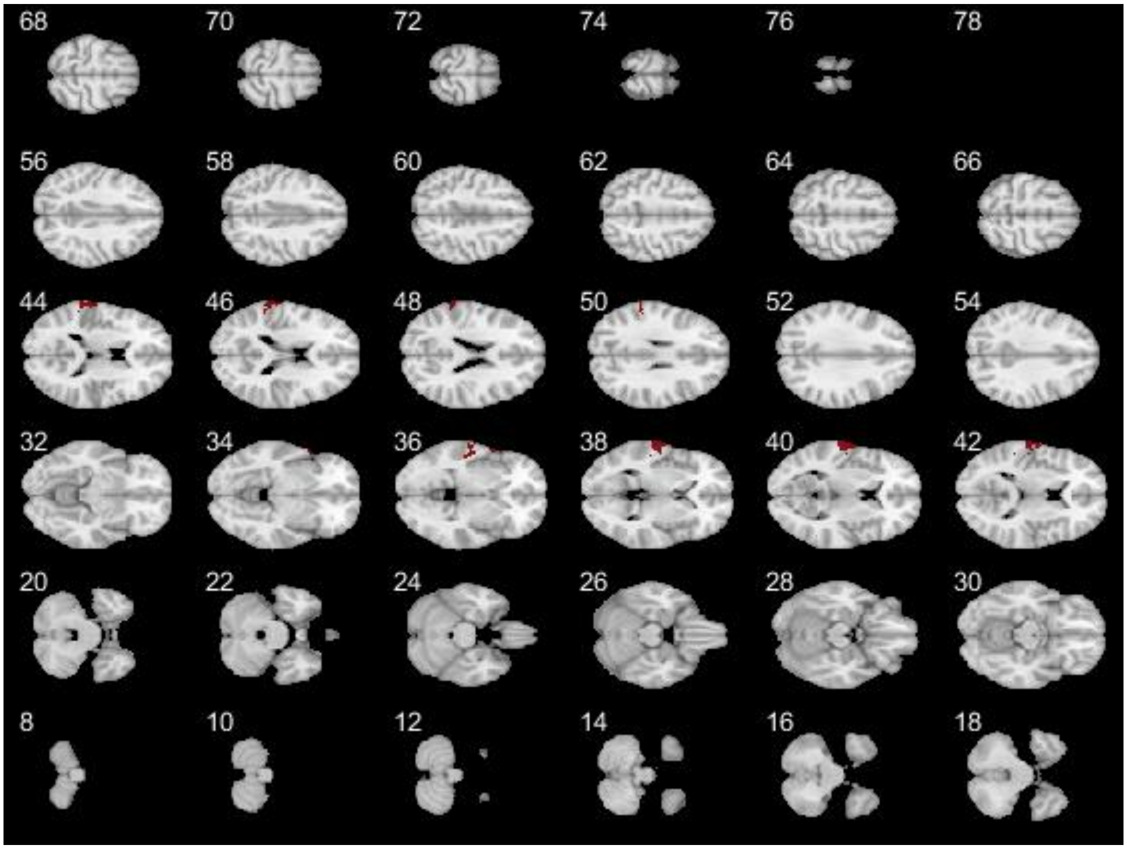


Figure S6: Significant cluster for Openness & Happy music using the ROV analysis. This result was found with a CDT of $p$ = 0.05, corrected at 5% FWER using non-parametric cluster-size and -mass thresholds, which were $k$ = 507 voxels and Z = 1295.28, respectively.

#### Discussion

##### Neuroticism & sad music

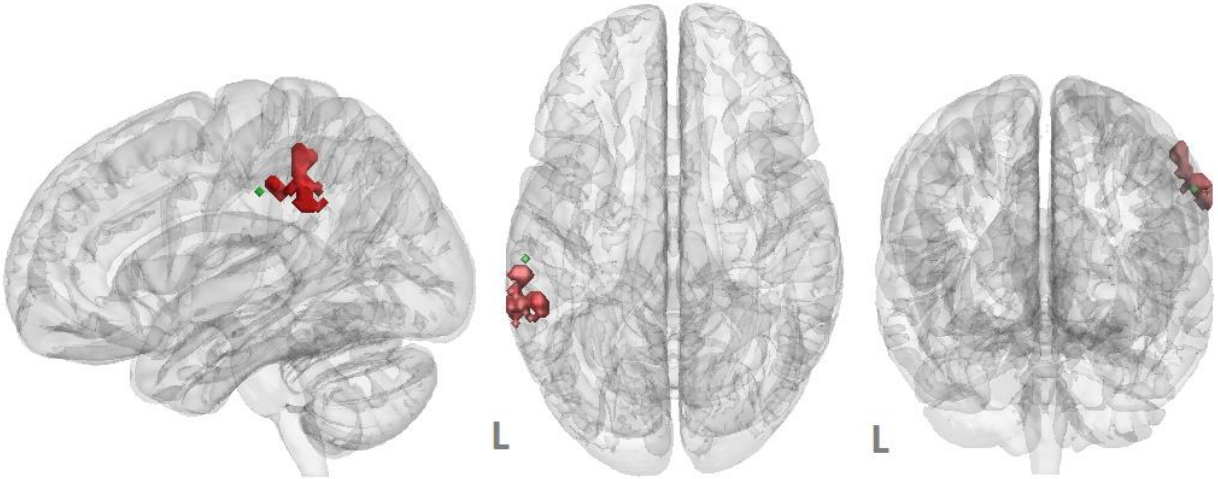


Figure S7: Peak voxel for the left IPL cluster in the action-observation network according to the meta-analysis by Caspers et al. (2010) (green), and the current results for the ROV analysis of Neuroticism and sad music (red).

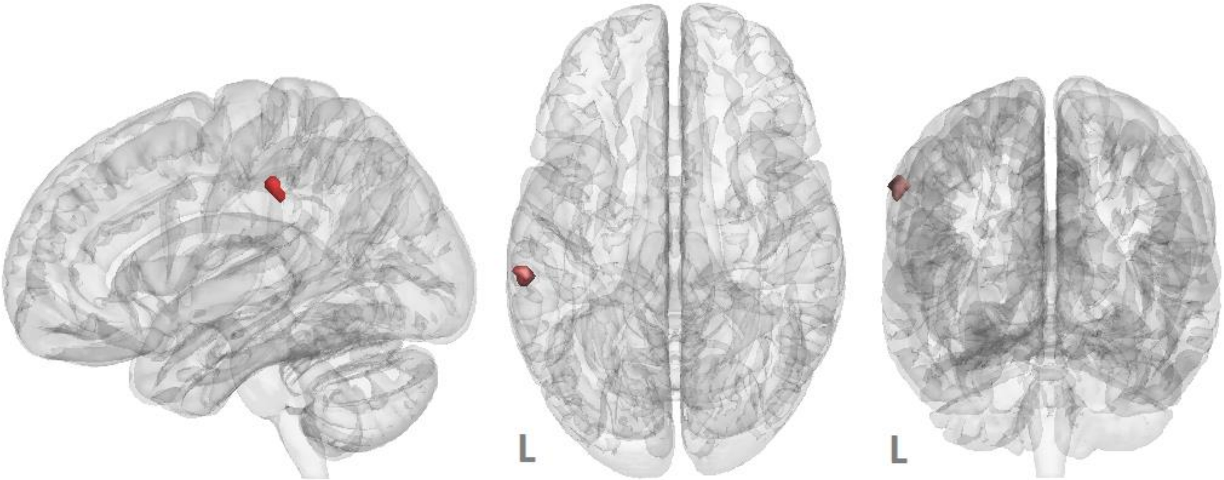


Figure S8: Intersection between the ROV results for Neuroticism and Sad music and association map for the term-based meta-analysis on the term “action observation” from NeuroSynth (Yarkoni et al., 2011; performed on March 29, 2019).

### References

Caspers, S., Zilles, K., Laird, A. R., & Eickhoff, S. B. (2010). ALE meta-analysis of action observation and imitation in the human brain. *NeuroImage, 50* (3), 1148–1167.

Dwass, M. (1957). Modified randomization tests for nonparametric hypotheses. *The Annals of Mathematical Statistics*, 181–187.

Edgington, E. S. (1969). Approximate randomization tests. *The Journal of Psychology*,

*72* (2), 143–149.

Kriegeskorte, N., Simmons, W. K., Bellgowan, P. S., & Baker, C. I. (2009). Circular analysis in systems neuroscience: The dangers of double dipping. *Nature Neuroscience*, *12* (5), 535.

Nichols, T. E., & Holmes, A. P. (2002). Nonparametric permutation tests for functional neuroimaging: A primer with examples. *Human Brain Mapping*, *15* (1), 1–25.

Omura, K., Aron, A., & Canli, T. (2005). Variance maps as a novel tool for localizing regions of interest in imaging studies of individual differences. *Cognitive, Affective, & Behavioral Neuroscience*, *5* (2), 252–261.

Suslow, T., Kugel, H., Lindner, C., Dannlowski, U., & Egloff, B. (2017, January). Brain response to masked and unmasked facial emotions as a function of implicit and explicit personality self-concept of extraversion. *Neuroscience*, *340* .

Yarkoni, T., Poldrack, R. A., Nichols, T. E., Van Essen, D. C., & Wager, T. D. (2011). Large-scale automated synthesis of human functional neuroimaging data. *Nature Methods*, *8* (8), 665.
